# Visualizing cancer-originated acetate uptake through MCT1 in reactive astrocytes demarcates tumor border and extends survival in glioblastoma patients

**DOI:** 10.1101/2021.04.13.439750

**Authors:** Hae Young Ko, Jee-In Chung, Dongwoo Kim, Yongmin Mason Park, Han Hee Jo, Sangwon Lee, Seon Yoo Kim, Jisu Kim, Joong-Hyun Chun, Kyung-Seok Han, Misu Lee, Yeonha Ju, Sun Jun Park, Ki Duk Park, Min-Ho Nam, Youngjoo Park, Se Hoon Kim, Jin-Kyoung Shim, Seok-Gu Kang, Jong Hee Chang, C. Justin Lee, Mijin Yun

**Affiliations:** Department of Nuclear Medicine, Severance Hospital, Yonsei University College of Medicine, Seoul 03722, Republic of Korea; CONNECT-AI Research Center, Yonsei University College of Medicine, Seoul 03722, Republic of Korea; Center for Cognition and Sociality, Institute for Basic Science, Daejeon 34126, Republic of Korea; IBS School, University of Science and Technology, Daejeon 34126, Republic of Korea; Department of Biological Sciences, Chungnam National University, Daejeon 34134, Republic of Korea; Division of Life Science, College of Life Science and Bioengineering, Incheon National University, Incheon 22012, Republic of Korea; Convergence Research Center for Diagnosis, Treatment and Care System of Dementia, Korea Institute of Science and Technology (KIST), Seoul 02792, Republic of Korea; Division of Bio-Med Science & Technology, KIST School, Korea University of Science and Technology, Seoul 02792, Republic of Korea; Center for Neuroscience, KIST, Seoul 02792, Republic of Korea; Department of KHU-KIST Convergence Science and Technology, Kyung Hee University, Seoul 02447, Republic of Korea; Department of Pathology, Severance Hospital, Yonsei University College of Medicine, Seoul 03722, Republic of Korea; Department of Neurosurgery, Severance Hospital, Yonsei University College of Medicine, Seoul 03722, Republic of Korea; Yonsei University College of Medicine, Seoul 03722, Republic of Korea

**Keywords:** Acetate, Glioblastoma, Image-guided surgery, PET image, Reactive astrocytes

## Abstract

Glioblastoma multiforme (GBM) is a devastating brain tumor with dismal prognosis of only 15-month survival regardless of surgical resection. Here, we report an advanced neuroimaging technique combining ^11^C-acetate PET and MRI (AcePET), visualizing the boundary beyond the MRI-defined tumor. Targeted biopsy of the regions with increased ^11^C-acetate uptake revealed the presence of reactive astrocytes with enhanced acetate-transporter MCT1, along with cancer stem cells. Reactive astrogliosis and MCT1-dependent ^11^C-acetate-uptake were recapitulated in U87MG-orthotopic models. Mechanistically, glycolytic tumor cells release excessive acetate causing reactive astrogliosis, leading to the release of aberrant astrocytic GABA and H_2_O_2_, which further down-regulate the neuronal glucose uptake through GLUT3. Clincally, AcePET-guided surgery allows complete tumor resection of infiltrating cancer stem cells and extends the overall survival of patients by 5.25 months compared to conventional MRI-guided surgery. We established a new concept of the metabolic interactions between GBM cells and neighboring neurons through reactive astrocytes and developed AcePET-guided surgery to fight against GBM.

## Introduction

Glioblastoma multiforme (GBM), the most aggressive type of glioma with a median overall survival (OS) of about 15 months, accounts for the majority of primary malignant tumors in the brain (Van Meir et al., 2010). Currently, the standard treatment modality is surgical resection, where the margins are determined by either magnetic resonance imaging (MRI) or 5-aminolevulinic acid (5-ALA) fluorescence. The strongest evidence for the use of T1 gadolinium-enhanced MRI comes from a randomized controlled trial: A complete resection of contrast-enhanced tumor resulted in longer OS compared to an incomplete resection (Pichlmeier et al., 2008). However, regardless of the complete surgical resection based on MRI, recurrence happens in almost all cases mostly at the margin of resection cavity in the peri-tumoral area (Petrecca et al., 2013). Therefore, the current MRI-guided method of surgery has a limitation of not detecting the true tumor margin.

As the latest technology, 5-ALA fluorescence-guided surgery has been helpful for surgeons to identify the apparent tumor margin intraoperatively (Stummer et al., 2000). A meta-analysis yielded a pooled OS difference of 3.05 months, favouring 5-ALA-guided surgery compared with control groups without 5-ALA (Gandhi et al., 2019). However, one of the major limitations of 5-ALA-guided surgery is the vague boundary fluorescence due to insufficient 5-ALA uptake in low tumor-cell density at the advancing tumor margin (Utsuki et al., 2006). To make matters worse, false-positive fluorescence is common due to the leakage of fluorescent molecules into the extracellular matrix (Utsuki et al., 2007). Despite the assistance of 5-ALA fluorescence, it is still difficult to achieve a complete resection of scattered cancer cells in the peri-tumoral region. Therefore, there is a desparate need for advanced imaging techniques that clearly visualize the peri-tumoral infiltrating cancer cells, allowing for more complete resection and extended patient survival.

MRI and 5-ALA fluorescence do not account for the presence of cancer stem cells and other non-tumorous cells such as microglia/macrophages, astrocytes, neutrophils, dendritic cells, endothelial cells in the tumor microenvironment (TME) (Quail and Joyce, 2017). Astrocytes undergo the process of reactive astrogliosis by changing their transcriptional profile and morphology, which is characterized by up-regulation of glial fibrillary acidic protein (GFAP) (Pekny and Nilsson, 2005). We have recently demonstrated that reactive astrocytes, especially scar-forming severe reactive astrocytes, generate excessive GABA and H2O2 via MAO-B-dependent putrescine degradation pathway causing neuronal dysfunction and neurodegeneration in Alzheimer’s disease (Chun et al., 2020; Jo et al., 2014). In the TME, reactive astrocytes are thought to enhance the proliferation, invasion, and drug resistance of GBM by secreting cytokines and chemokines (Le et al., 2003; Lee et al., 2011; Nagashima et al., 2002; Placone et al., 2016). Specifically, they up-regulate tumor cell proliferation by secreting growth factors such as IL-6, TGF-β and IGF-1 (Placone et al., 2016) and influence the invasion of tumor cells by the cleavage of inactive pro-MMP2 to active MMP2 (Le et al., 2003). In the metastatic brain tumor model, reactive astrogliosis and metastatic tumor cells stimulate each other and this mutual relationship contributes to the development of metastasis in the brain (Seike et al., 2011). Reactive astrocytes are shown to surround glioma in both human biopsy tissues (Nagashima et al., 2002) and murine models (Lee et al., 2011). Other than tumor initiation and progression, reactive astrocytes remaining after resection promote tumor proliferation and invasion in a murine glioma resection and recurrence model (Okolie et al., 2016), raising the possibility that cancer stem cells are present in regions of reactive astrogliosis. Thus, it would be highly advantageous to develop a functional imaging technique to visualize reactive astrogliosis *in vivo* and aid image-guided complete tumor resection. However, there has yet to be an attempt to specifically visualize reactive astrocytes surrounding GBM in clinics.

^11^C–acetate, a radiotracer version of acetate, is currently in clinical use to detect low-grade tumors with acetate dependence in the body (Yun et al., 2009). In the brain, we unexpectedly discovered that high-grade gliomas were associated with high ^11^C-acetate uptake in patients on positron emission tomography (PET) (Kim et al., 2018). It has been recently reported that acetate is a bioenergetic substrate for human GBM (Mashimo et al., 2014), alluding to the possibility that GBM cells take up acetate. In contrast, few prior clinical studies on patients with multiple sclerosis reported that reactive astrocytes are presumably responsible for increased ^11^C–acetate uptake (Kato et al., 2021). Such a discrepancy raises the question on which cells in high grade gliomas are taking up ^11^C-acetate. In this study, we investigated whether reactive astrocytes surrounding gliomas take up high levels of ^11^C-acetate, delineated the molecular mechanisms of acetate-dependent reactive astrogliosis, and assessed the OS of patients with high-grade gliomas who underwent ^11^C-acetate PET and MRI (AcePET) guided surgery.

## Results

### Reactive astrogliosis at the tumor boundary in patients with high-grade gliomas

To test the possibility that reactive astrocytes surrounding gliomas take up high levels of ^11^C-acetate, we compared tumor volumes between ^11^C-acetate PET and MRI images in IDH1-WT high-grade gliomas (Figure 1A), which provided the basis for AcePET-guided surgery (Figure 1B). On visual inspection, IDH1-WT high-grade gliomas showed significantly higher ^11^C-acetate uptake at the periphery of the tumor (Figures 1A; red demarcation and 1C), while low-grade or IDH1-mt tumors demonstrated minimal or no significant ^11^C-acetate uptake. The median standard uptake value (SUV) ratio (SUVR) on ^11^C-acetate PET in IDH1-WT high-grade glioma (*n*=37) was significantly increased compared to that in low-grade or IDH1-mt glioma (*n*=58) (2.05 (Interquartile range (IQR) 1.77-2.41) versus 1.02 (IQR 0.92-1.30)). The calculated tumor volume by ^11^C-acetate PET was remarkably larger than that of contrast-enhanced MRI (Figure 1A, blue demarcation) in IDH1-WT high-grade gliomas (Figures 1C and 1D). These results indicate that ^11^C-acetate PET exhibits a clear and larger tumor boundary beyond the contrast-enhanced MRI.

**Figure 1.**
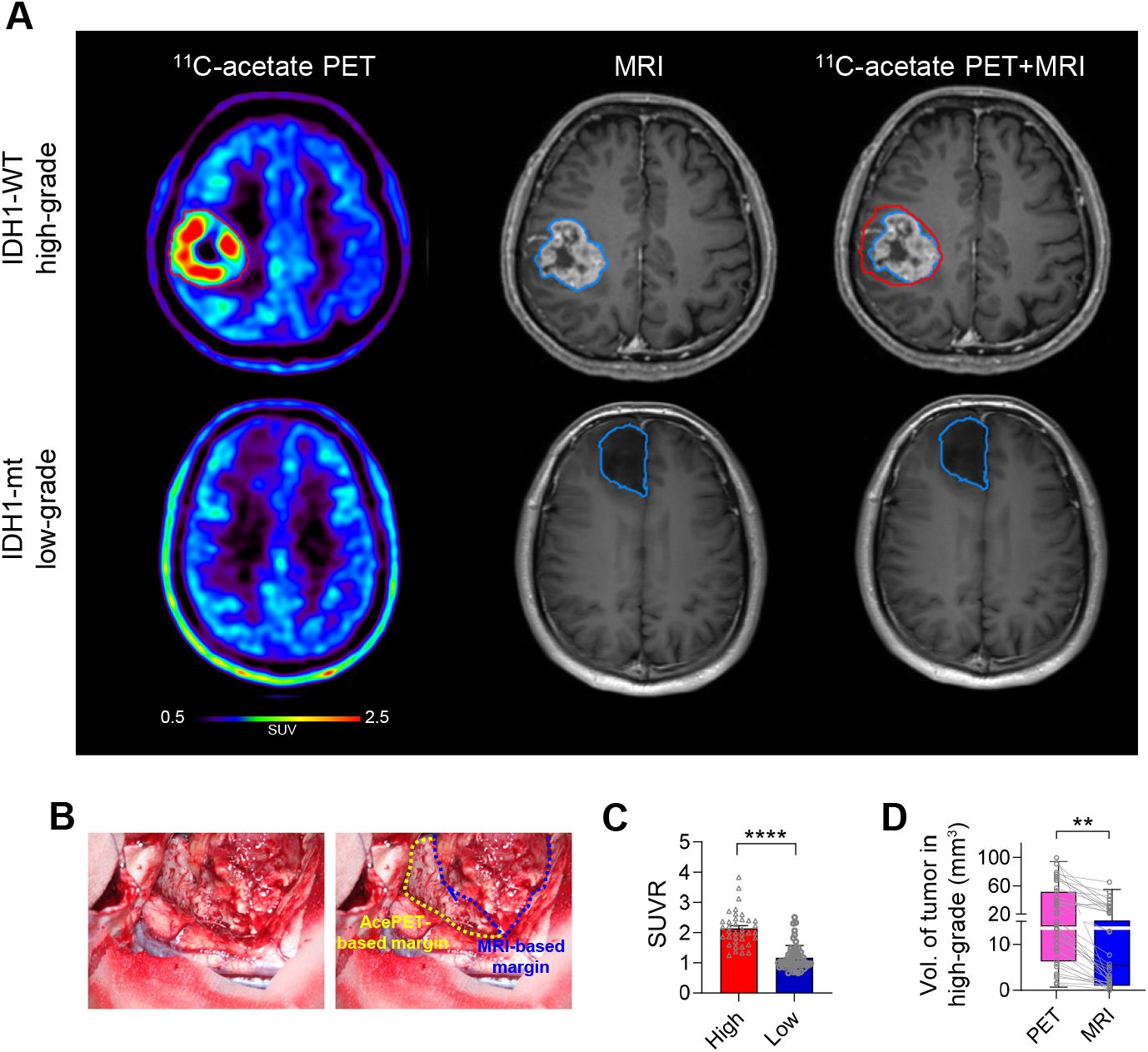
(A) Representative images of ^11^C-acetate PET, MRI, and ^11^C-acetate PET-MRI fusion in patients with IDH1-WT high-grade and IDH1-mt low-grade glioma. Red demarcation, ^11^C-acetate PET-based tumor margin; Blue demarcation, MRI-based tumor margin. (B) Intraoperative images of IDH1-WT high-grade glioma with AcePET (yellow)- or MRI (blue)-based margins. (C) The median SUVR on ^11^C-acetate PET in IDH1-WT high-grade glioma (*n*=37) was significantly increased than that in low-grade or IDH1-mt glioma (*n*=58) (2.05 (IQR 1.77-2.41) versus 1.02 (IQR 0.92-1.30)). (D) The median tumor volume on ^11^C-acetate PET was significantly larger than that on MRI in IDH1-WT high-grade gliomas (*n*=37) (20.80 (IQR 8.99-59.39) versus 7.72 (IQR 1.80-26.14)). Data are presented as median and IQR by Mann-Whitney U-test (C) and unpaired two-tailed t-test (D). **P < 0.01, ****P < 0.0001.

To investigate whether reactive astrogliosis is present at high ^11^C-acetate uptake-regions, we performed double-immunohistochemistry with targeted biopsy tissues of high- and low-grade gliomas using antibodies against GFAP, an astrocyte marker, and Ki-67, a marker for proliferating cells (Figures 2A and 2B). In IDH1-WT high-grade glioma, we identified three distinct regions by the presence of tumor and GFAP-positive astrocytic morphology; the T tumor region with amoeboid-shaped astrocytes, the L region with line-shaped astrocytes, and the S region with star-shaped astrocytes (Figures 2A, 2B, S1A and S1B). The tangential line-shaped astrocytes (Figure 2B) delineated the boundary between T and S regions to form a glial scar-like barrier around the tumor. In contrast to IDH1-WT high-grade glioma, we found no T or L regions and only S regions in low-grade or IDH1-mt gliomas (Figures 2B and 2D). Astrocyte reactivity at the margin of gliomas was more severe in highgrade gliomas than in low-grade or IDH1-mt gliomas, as proven by higher expressions of both GFAP and MAO-B, a recently identified reactive astrocyte marker (Jo et al., 2014) (Figures 2C, 2D, 2G and 2I). MCT1, a potential acetate transporter in astrocytes (Jeon et al., 2018; Waniewski and Martin, 1998), was found to be expressed at high levels in the periphery of IDH1-WT high-grade gliomas, but almost absent in low-grade or IDH1-mt gliomas (Figures 2C, 2D and 2J), which was consistent with the ^11^C-acetate uptake results (Figures 1A and 1C). These results indicate that reactive astrogliosis is present at high ^11^C-acetate uptake-regions, particularly at the tumor margin of high-grade gliomas. Moreover, we found numerous Ki-67-positive cells in L and S regions of high-grade gliomas (Figures 2A, 2C and 2H), but not in low-grade or IDH1-mt gliomas (Figures 2B, 2D and 2H). The Ki-67-positive cells were mostly co-localized with CD133, the marker for cancer stem cells (Liu et al., 2006) (Figures 2E and 2K), proving the existence of cancer stem cells within reactive astrogliosis at the tumor margin. We also found MCT1 expression in CD133-positive cancer stem cells in the tumor margin (Figure 2F), which were very low in number and intensity compared to reactive astrocytes (7.24% in L region; 8.39% in S region) (Figure 2L).

**Figure 2.**
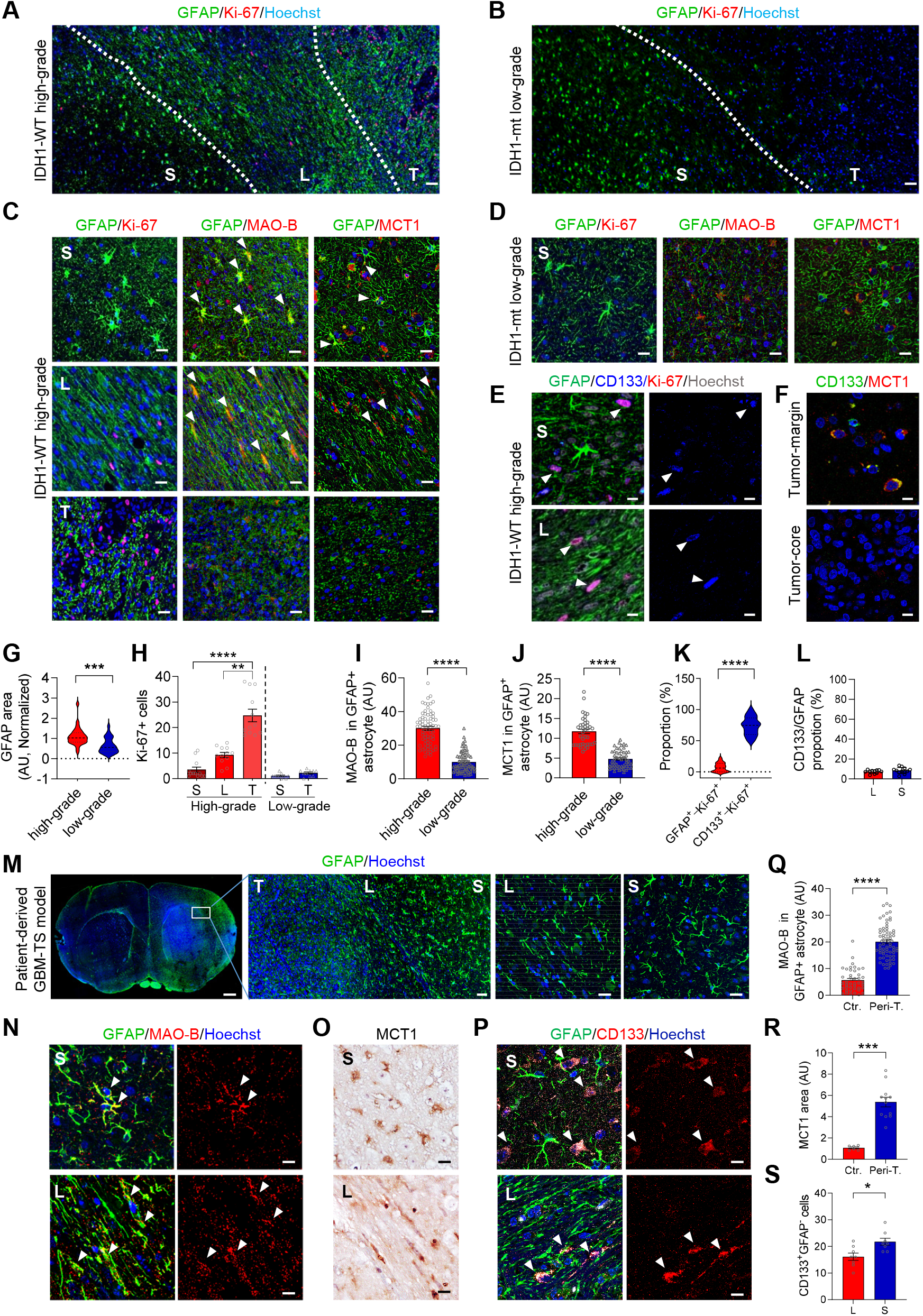
Reactive astrogliosis resides in the peri-tumoral region on patients with high-grade gliomas and GBM-TS models. (A-D) Immunofluorescence images with GFAP, Ki-67, MAO-B and MCT1 in human glioma tissues. S, Star-shape astrocyte region; L, Linear-shape astrocyte region; T, Tumor region. (E) Immunofluorescence images with GFAP, Ki-67, and CD133 in the peri-tumoral region on IDH1-WT high-grade glioma tissues. (F) Immunofluorescence images with CD133 and MCT1 in the peri-tumoral region and tumor core region on IDH1-WT high-grade glioma tissues. (G-I) Quantification of GFAP, Ki-67 and MAO-B immunoreactivity in IDH1-WT high-grade versus IDH1-mt or low-grade gliomas (*n*=3). (J) Quantification of MCT1 immunoreactivity in GFAP+ astrocyte or CD133+ cells. (*n*=3) (K) The proportion of GFAP+ or CD133+ in Ki-67+ cells in IDH1-WT high-grade glioma tissues (*n*=4). (L) The proportion of CD133 versus GFAP in L and S region. (M-O) Immunofluorescence images with GFAP, MAO-B and MCT1 in patient-derived GBM-TS models. (P) Immunofluorescence images with GFAP and CD133 in GBM-TS models. (Q-R) Quantification of MAO-B and MCT1 immunoreactivity on the peri-tumoral (Peri-T.) and contralateral regions (Ctr.) in GBM-TS models (*n*=3). (S) Quantification of CD133+ and GFAP-cells in L and S region. Scale bars, 50 μm in (A, B, and M middle); 20 μm in (C, D, and M right); 10 μm in (E, F, N, O, and P); 500 μm in (M left). Data are presented as mean ± SEM. *P < 0.05, **P < 0.01, ***P <0.001, ****P < 0.0001 by unpaired two-tailed t-test (G, I, K, Q, R and S), Kruscal-wallis test (H left), Mann-Whitney U-test (H right) or one way ANOVA with Turkey (J).

To investigate whether the presence of reactive astrogliosis and cancer stem cells can be recapitulated in a mouse orthotopic xenograft model, we developed patient-derived xenografts using tumorsphere isolated from high-grade glioma patients (GBM-TS model; Figure 2M) as previously described (Jeong et al., 2019; Park et al., 2018). We then performed immunostaining with antibodies against GFAP, MAO-B, MCT1 and CD133 (Figures 2M, 2N, 2O and 2P) and found the presence of distinct T, L and S regions with GFAP-positive reactive astrocytes at the tumor margin (Figure 2M). Similar to high-grade gliomas, these reactive astrocytes displayed strong expressions of GFAP, MAO-B and MCT1 at peri-tumoral L and S regions compared to contralateral regions (Figures 2N, 2O, 2Q and 2R). In addition, we found CD133-postive cancer stem cells, scattered within the region of reactive astrogliosis in the peri-tumoral region (Figures 2P and 2S). These results indicate that reactive astrogliosis and cancer stem cells exist around the tumor in patient-derived GBM-TS models.

### Inhibition of MAO-B and MCT1 decreases reactive astrogliosis and ^11^C-acetate uptake

To delineate the molecular and cellular mechanism of elevated ^11^C-acetate uptake in the peri-tumoral region of high-grade glioma, we employed a reverse-translational approach and performed ^11^C-acetate microPET imaging and ^14^C-acetate autoradiography in orthotopic mouse tumor models. We used two types of human glioma cell lines; U87MG cell line to mimic high-grade glioma and U87-IDH1-mt (U87mt) to mimic IDH1-mt glioma. We placed a region-of-interest (ROI) over the peri-tumoral and mirror-image contralateral regions to measure the SUV ratio on ^11^C-acetate microPET images. We found a significantly higher SUV ratio in the peri-tumoral region in the U87MG model compared to the U87mt model (Figures 3A and 3E). We then asked whether the elevated ^11^C-acetate uptake is associated with reactive astrogliosis and astrocytic MCT1 level. To detect the presence of reactive astrogliosis in U87MG and U87mt models, we immunostained with antibodies against GFAP, S100β (another reactive astrocyte marker), MAO-B, and MCT1(Figures 3B, 3C and 3D). Similar to human gliomas, the U87MG model showed significantly higher intensity and a larger area of GFAP and S100β immunoreactivities than the U87mt model (Figures 3F 3G and 3H). MAO-B and MCT1 levels in GFAP- and S100β-positive areas in the peri-tumoral region were also significantly higher in U87MG than in U87mt models (Figures 3C, 3D, 3I and 3J). These results indicate that ^11^C-acetate uptake was positively correlated with both reactive astrogliosis and astrocytic MCT1.

**Figure 3.**
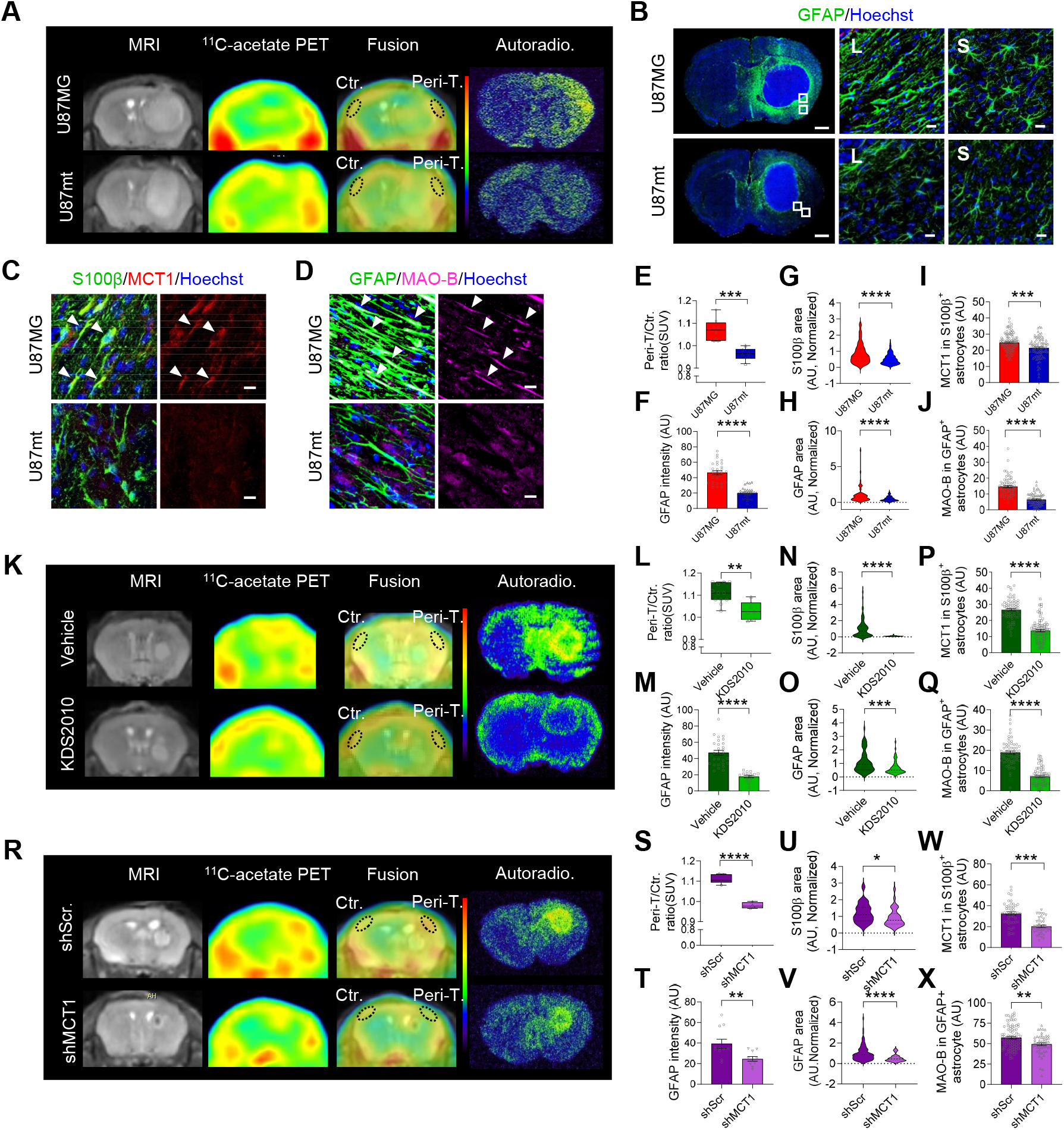
Inhibition of MAO-B and MCT1 decreases reactive astrogliosis and acetate uptake in mouse glioblastoma model. (A) Representative images of MRI, ^11^C-acetate PET, PET and MRI fusion, and ^14^C-acetate autoradiography in mouse glioma models of U87MG and U87mt. (B) Immunofluorescence images of GFAP expression in the mouse models. (C-D) Immunofluorescence images of S100β and MCT1 expressions, GFAP and MAO-B expressions in the mouse models. (E) SUVR of ^11^C-acetate in the peri-tumoral and contralateral regions (*n*=5). (F-J) Quantification of GFAP, S100β, MCT1, and MAO-B expressions in the mouse models (*n*=3). (K) Representative images of MRI, ^11^C-acetate PET, PET and MRI fusion, and ^14^C-acetate autoradiography with or without KDS2010 administration in the U87MG mouse model. (L) SUVR of ^11^C-acetate in the peri-tumoral (Peri-T.) and contralateral regions (Ctr.) (*n*=5). (M-Q) Quantification of GFAP, S100β, MCT1, and MAO-B expressions with or without KDS2010 administration in U87MG mouse models (*n*=3). (R) Representative images of MRI, ^11^C-acetate PET, PET and MRI fusion and ^14^C-acetate autoradiography with or without astrocyte-specific MCT1 gene-silencing around U87MG cells. (S) SUVR of ^11^C-acetate in the peri-tumoral and contralateral regions (*n*=5). (T-X) Quantification of GFAP, S100β, MCT1, and MAO-B expressions with or without astrocyte-specific MCT1 gene-silencing of around U87MG cells (*n*=3). Scale bars, 500 μm in (B left); 10 μm in (B right, C, and D), Data are presented as mean ± SEM. *P < 0.05, **P < 0.01, ***P < 0.001, ****P < 0.0001 by unpaired two-tailed t-test (E, F, L, M, S, T, W, and X) or Mann-Whitney U-test (G-J, N-Q, U, and V).

To test whether reactive astrogliosis is necessary for the elevated ^11^C-acetate uptake in the U87MG model, we used the selective and reversible MAO-B inhibitor KDS2010, which has been shown to block reactive astrogliosis in animal models of Alzheimer’s disease, Parkinson’s disease, and subcortical stroke in our previous reports (Heo et al., 2020; Jo et al., 2014; Nam et al., 2020; Park et al., 2019). After oral administration of KDS2010, we found significantly decreased ^11^C-acetate uptake on microPET images and decreased ^14^C-acetate uptake on autoradiography at the peri-tumoral regions (Figures 3K and 3L). In addition, GFAP, S100β, MAO-B, and MCT1 levels in reactive astrocytes were all significantly reduced after KDS2010 treatment (Figures 3M–3Q, S2A and S2B). These results indicate that reactive astrogliosis is necessary for elevated ^11^C-acetate uptake.

To test whether astrocytic MCT1 is necessary for ^11^C-acetate uptake, we adopted a cre-loxp-dependent, astrocyte-specific gene-silencing of MCT1 using AAV-GFAP-cre-mCherry and AAV-pSico-rMCT1sh-GFP viruses around the tumor in the U87MG model. We found that gene-silencing of MCT1 significantly decreased ^11^C-acetate uptake on microPET images and ^14^C-acetate uptake on autoradiography at the peri-tumoral regions (Figures 3R and 3S). In addition, the levels of GFAP, S100β, and MAO-B were significantly reduced by the astrocytic gene-silencing of MCT1 (Figures 3T–3X, S2C and S2D), indicating that astrocytic MCT1 is necessary for elevated ^11^C-acetate uptake. Taken together, the elevated ^11^C-acetate uptake requires both reactive astrogliosis and astrocytic MCT1 at the boundary of high-grade tumor.

### Molecular factors of high-grade glioma that induce reactive astrogliosis

To delineate the detailed molecular and cellular mechanisms behind the elevated acetate uptake, we performed various *in vitro* biochemical and physiological assays with U87MG and mouse primary astrocytes (AST). We first screened for the cell types that show a high level of MCT1. We found that AST showed much higher expression of MCT1 along with GFAP and MAO-B compared to various glioma cell lines, including U87MG, U373MG, T98G, and U87mt (Figure 4A). Consistently, AST showed much higher ^14^C-acetate uptake compared to the aforementioned glioma cell lines (Figure 4B). These results implicate that the cellular origin of the enhanced acetate uptake is most likely attributed to astrocytes rather than tumor cells. To identify the tumor-derived molecular factors that induce reactive astrogliosis, we obtained conditioned media (CM) from U87MG or U87mt cells and applied it to AST (Figure 4C). U87MG-CM, but not U87mt-CM, significantly augmented MAO-B and MCT1 expression (Figure 4D) and ^14^C-acetate uptake (Figure 4E). U87MG-CM also increased mRNA levels of MCT1(Figure 4F). Gene-silencing of MCT1 in AST (Figures 4G and H) significantly prevented ^14^C-acetate uptake (Figure 4I), indicating that MCT1 is necessary for acetate uptake. These results indicate that the tumor-derived molecular factors induced reactive astrogliosis with the elevated expression of MAO-B and MCT1.

**Figure 4.**
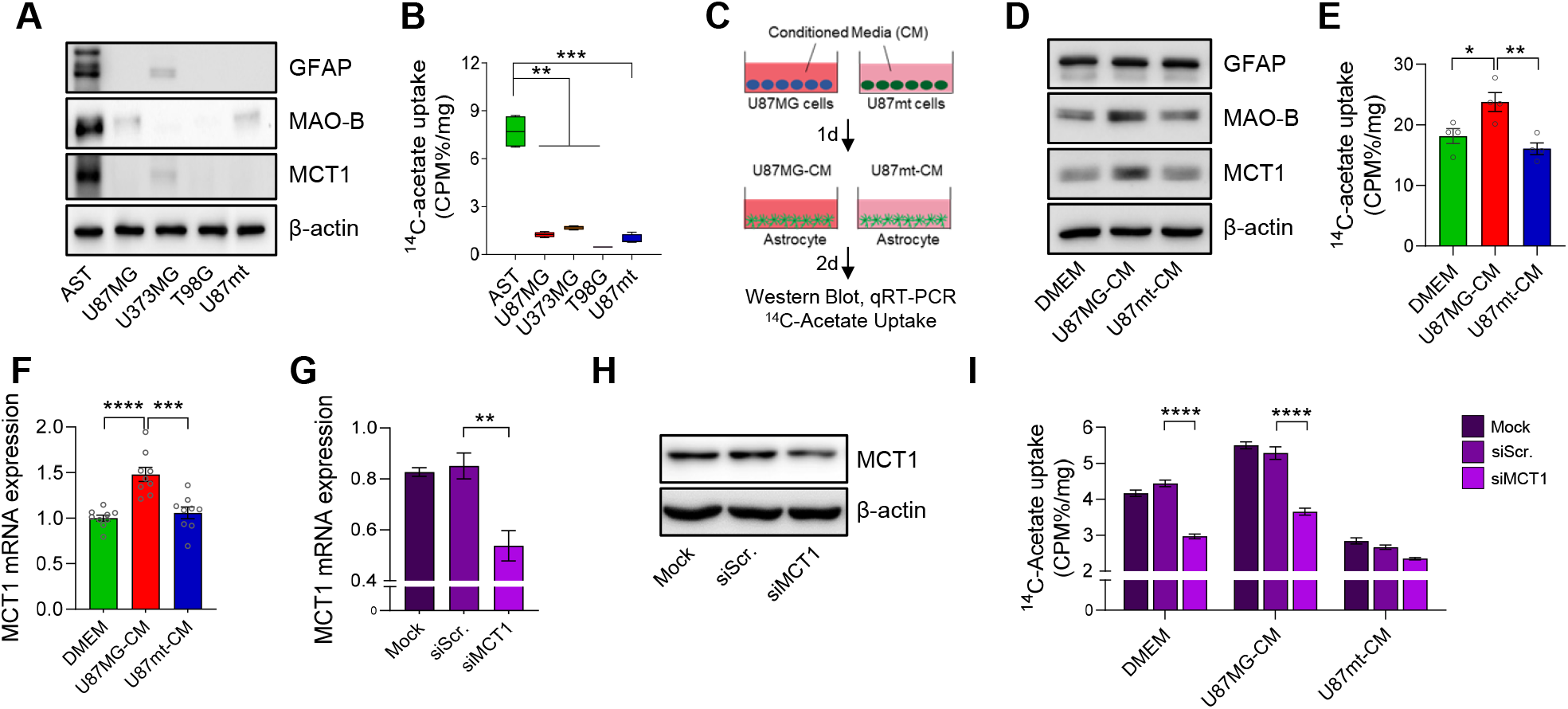
Excessive acetate from high-grade glioma is taken up to cause reactive astrogliosis. (A-B) Western-blot for GFAP, MAO-B, and MCT1 expressions and ^14^C-acetate uptake in mouse primary astrocytes (AST) and various human GBM cell lines (U87MG, U373MG, T98G, and U87mt). (C) Schematic diagram of preparing U87MG or U87-IDH1mutant-Conditioned Media (CM). (D) Western-blot for GFAP, MAO-B, and MCT1 expressions. (E) ^14^C-acetate uptake (F) MCT1 mRNA expressions in U87MG and U87mt-CM-treated astrocytes. (G-H) MCT1 expression (mRNA and protein) in siMCT1-transfected AST. (I) ^14^C-acetated uptake after siMCT1-transfection in U87MG-CM and U87mt-CM-treated astrocytes. Data are presented as mean ±SEM. *P < 0.05, **P < 0.01, ***P < 0.001, ****P < 0.0001 by one-way ANOVA with Tukey (B, E, F, G, and I)

### Excessive acetate from high-grade glioma is taken up to cause astrocyte reactivity

Based on the fact that reactive astrocytes take up high levels of ^14^C-acetate, we hypothesized that acetate is one of the tumor-derived metabolites that cause reactive astrogliosis. To verify this hypothesis, we measured the amount of acetate in U87MG-CM using an aceate assay kit (Figure 5A). We found a time-dependent increase of secreted acetate from U87MG (Figure 5A), consistent with a previous report on colorectal cancer (Liu et al., 2018). To investigate whether acetate alone induces reactive astrogliosis, we treated AST with various concentrations of acetate and found that aceate increased expression of MAO-B and MCT1 (Figure 5B). To determine whether MCT1 is responsible for reactive astrogliosis, we treated AST with AAV-GFAP-cre-mCherry and AAV-pSico-rMCT1sh-GFP viruses (Figure 5C). We found that gene-silencing of MCT1 prevented the acetate-induced increase of MAO-B-expression (Figure 5C).

**Figure 5.**
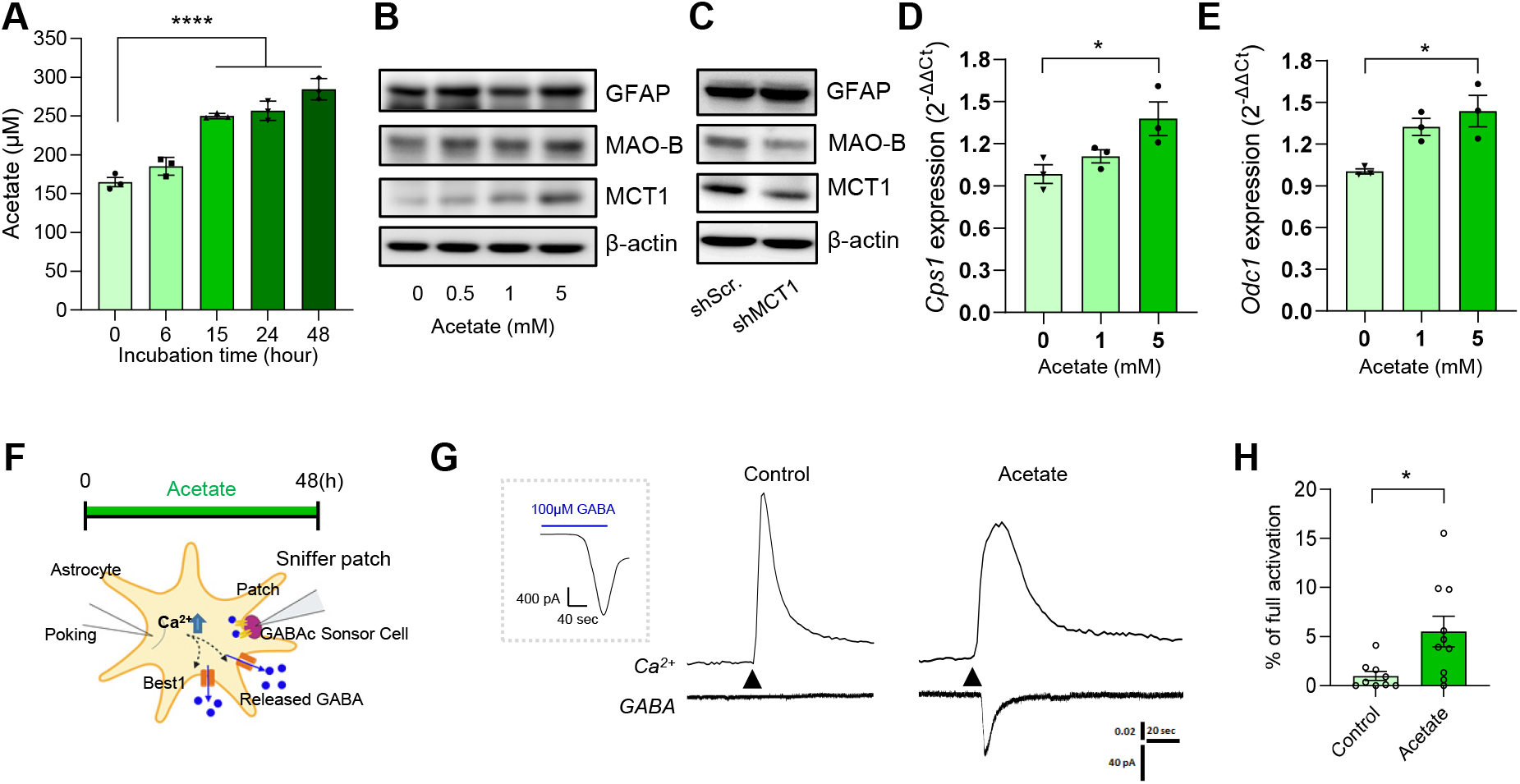
Excessive acetate from high-grade glioma causes reactive astrogliosis. (A) Concentration of acetate in U87MG-CM. (B) Western blot for GFAP, MAO-B and MCT1 expressions in AST treated with acetate. (C) Western blot for GFAP, MAO-B and MCT1 expressions in shMCT1-transfected AST. (D-E) mRNA expression of *Cps1* and *Odc1* in AST treated with acetate. (F) Schematic diagram of sniffer patch to record the astrocytic GABA release. (G) Representative traces of Ca^2+^ signal (top) and GABA current (bottom). Arrowheads indicate the time-point of poking the astrocyte. (H) Quantification of poking-induced GABA current in AST treated with acetate. Data are presented as mean ± SEM. *P < 0.05, ***P < 0.001 by one-way ANOVA with Tukey (A, D, and E) or unpaired two-tailed t-test (H).

We have recently demonstrated that putrescine, the source of astrocytic GABA, is increased via the urea cycle and polyamine metabolism in reactive astrocytes (Ju, 2021). To test whether acetate alone is associated with the urea cycle and polyamine metabolism, we measured the mRNA level of carbamoyl phosphate synthetase 1 (CPS1), the first enzyme of the urea cycle, and ornithine decarboxylase 1 (ODC1), the key enzyme behind polyamine biosynthesis converting ornithine into putrescine. We found that *Cps1* and *Odc1* mRNA levels were increased by acetate (Figures 5D and 5E), suggesting that it plays an important role in the urea cycle and polyamine metabolism, leading to GABA synthesis.

We have shown that tumor-derived acetate increased MAO-B expression in reactive astrocytes. This raises the interesting possibility that excessive acetate, after being taken up by astrocytes, can cause MAO-B-dependent GABA-synthesis and -release, which are closely associated with reactive astrogliosis (Chun et al., 2020; Chun and Lee, 2018; Jo et al., 2014). To test this possibility, we performed a sniffer-patch with GABAc-expressing HEK293T cells as a biosensor for GABA as previously described (Figure 5F) (Oh et al., 2019). We found that poking-induced, Ca^2+^-dependent GABA-release was significantly enhanced after a 2-day incubation with 5 mM acetate compared to the control (Figures 5G and 5H). These findings indicate that the excessive acetate causes reactive astrocytes, leading to MAO-B-dependent GABA-synthesis and -release, possibly via the Best1 channel.

### Astrocytic GABA and H2O2 induces neuronal glucose hypometabolism

What is the role of GABA from reactive astrocytes? We have previously shown that GABA from reactive astrocytes causes cortical glucose hypometabolism and impedes functional recovery after subcortical stroke (Nam et al., 2020). In addition, we have recently demonstrated that reactive astrogliosis-mediated glucose hypometabolism occurs via reduction in neuronal glucose transporter (GLUT3) in Alzheimer’s disease patients (Nam, 2021). To test if similar mechanisms exist in high-grade glioma, we measured the protein level of GLUT3 in *in vitro* neuron-glia co-culture, *in vivo* animal model, and glioma patients. In *in vitro* co-culture system, U87MG-CM treatment caused a significant decrease in the protein levels of neuronal nuclear protein (NeuN) and GLUT3, along with an increase in GFAP, MAO-B, and MCT1, in a time-dependent manner (Figure 6A). In *in vivo* animal models, we found a significantly higher level of astrocytic GABA (Figures 6B and 6J) in the peri-tumoral regions of the U87MG model than the U87mt model. Additionally, we found a significantly higher level of MAO-B-mediated H_2_O_2_ in U87MG-CM-treated astrocytes by DCFDA assay (Figure 6C). In primary cultured neurons, both GABA and H_2_O_2_ induced a significant decrease in ^14^C-deoxyglucose (^14^C-DG) uptake and the protein level of GLUT3 (Figures 6D and 6E). Consistently, we found lower levels of neuronal GLUT3 in the peri-tumoral region of the U87MG model compared to the U87mt model (Figures 6F and 6K).

**Figure 6.**
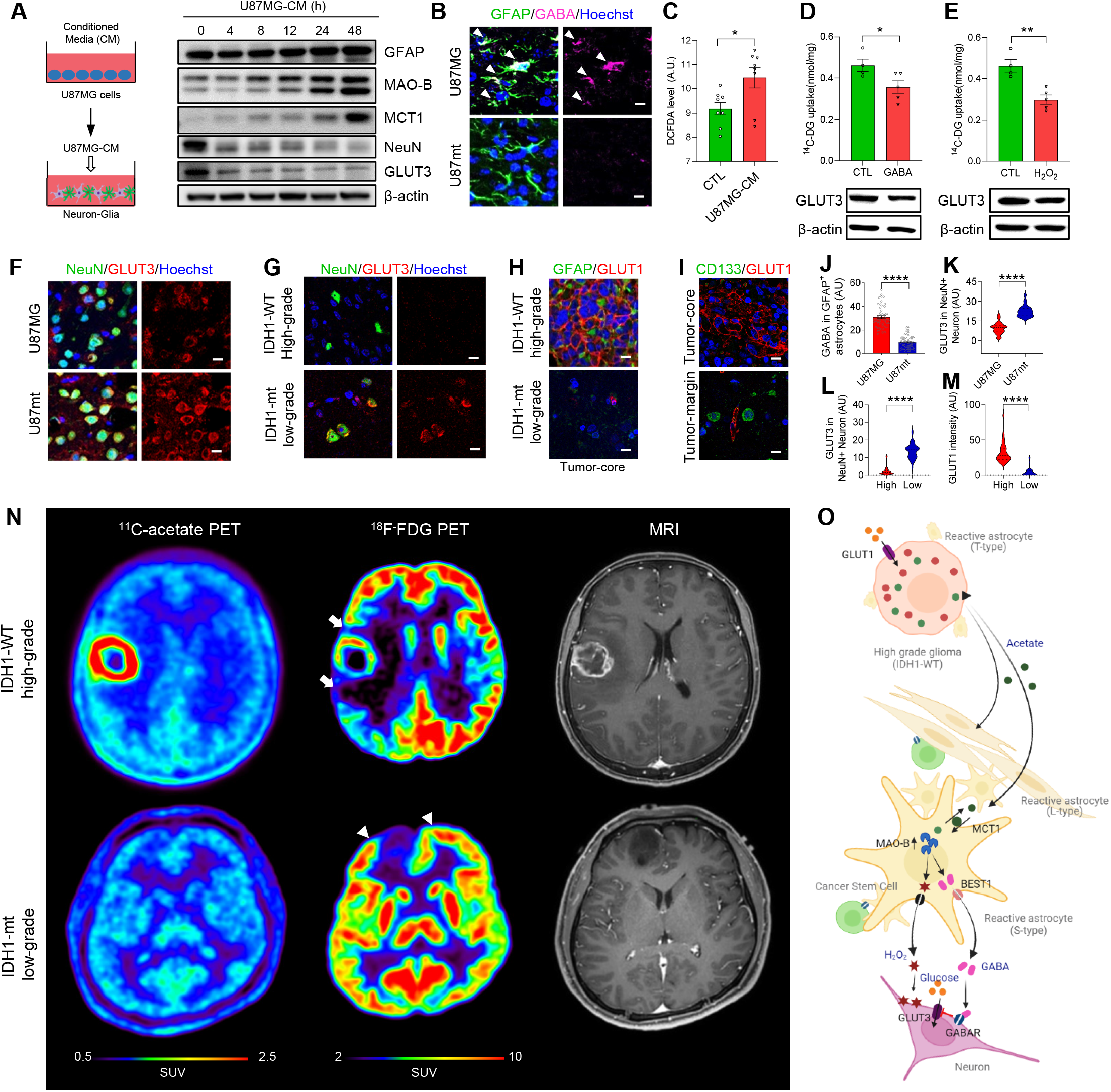
Astrocytic GABA suppresses neighboring neuronal glucose uptake in neuron-glia co-culture, mouse U87MG model, and patient high-grade gliomas. (A) Schematic diagram (left) and western-blot for GFAP, MAO-B, MCT1, NeuN, and GLUT3 from mouse primary neuron-glia co-culture treated with U87MG-CM. (B) Representative images of GFAP and GABA staining in U87MG and U87mt models. (C) DCFDA levels in U87-CM. (D-E) ^14^C-Deoxyglucose (DG) uptake (top) and western blot of GLUT3 (bottom) in primary neuron treated with GABA and H2O2. (F-G) Representative images of GLUT3 and NeuN staining in mouse models and human biopsy tissues. (H) Representative images of GFAP and GLUT1 staining in the tumor core region of human biopsy tissues. (I) Representative images of CD133 and GLUT1 staining in the tumor core and margin of high-grade glioma tissues. (J-K) Quantification of GABA in GFAP+ astrocytes and GLUT3 in NeuN+ neuron in mouse models (*n*=3). (L-M) Quantification of GLUT3 in NeuN+ neuron and GLUT1 expression levels in human biopsy tissues (*n*=3). (N) ^11^C-acetate PET, ^18^F-FDG PET, and MRI of patients with IDH1-WT high-grade glioma (top row) and IDH1-mt or low-grade glioma (bottom row). (O) Schematic illustration summarizing a hypothesized mechanism. Scale bars, 10 μm in (B, F, G, H, and I). Data are presented as mean ± SEM., *P < 0.05, **P < 0.01, ****P < 0.0001 by unpaired two-tailed t-test (C, D, E, J, and K) or Mann-Whitney U-test (L, and M).

In the targeted biopsy tissues, we found a significantly lower level of neuronal GLUT3 in IDH1-WT high-grade glioma than that of IDH1-mt or low-grade glioma (Figures 6G and 6L). In marked contrast, we observed a high level of GLUT1 in high-grade glioma, but not in IDH1-mt or low-grade glioma (Figures 6H and 6M). We also found a low level of GLUT1 in CD133-positive cancer stem cells in the tumor-margin (Figure 6I). In glioma patients, we found a remarkable decrease in ^18^F-fluorodeoxyglucose (FDG) uptake on PET images at the adjacent cortical regions beyond the tumor boundary, delineated by ^11^C-acetate PET in IDH1-WT high-grade glioma patients, but not in IDH1-mt or low-grade glioma patients (arrows and arrow heads in Figure 6N). These results indicate that the aberrant GABA and H_2_O_2_ from reactive astrocytes induce neuronal glucose hypometabolism. Taken together, these findings lead us to further postulate a multicellular mechanistic model, consisting of high-grade glioma cells that gorge on glucose and release excessive acetate, reactive astrocytes which release aberrant GABA and H_2_O_2_, and neighboring neurons with subsequently suppressed glucose metabolism (Figure 6O).

### AcePET-guided surgery extends OS in high-grade glioma patients

Based on the observation that cancer stem cells are present in regions of high ^11^C-acetate uptake at the tumor boundary beyond the contrast-enhanced MRI, we performed a Kaplan-Meier survival curve analysis to compare the OS in glioma patients who underwent conventional MRI-guided surgery versus AcePET-guided surgery. AcePET-guided surgery is performed under navigation by the fusion images of ^11^C-acetate PET and MRI, which were sent from the picture archiving and communication system (PACS) to the navigation system via local network (Figure S3A). There were 149 glioma patients with at least a 2-year follow-up after surgery who had received appropriate treatment according to the tumor grade (Figure S3B). The patients were divided into three groups depending on the tumor grade and the presence of the mutation p.R132H in isocitrate dehydrogenase 1 gene (IDH1-mt), which has been shown to exhibit a favorable disease outcome compared to its wild-type counterpart (IDH1-WT) (Louis et al., 2016); Group 1) Grade 2 gliomas (*n*=41), Group 2) IDH1-mt highgrade gliomas (*n*=28), and Group 3) IDH1-WT high-grade gliomas (*n*=80)(Figure S3B). The patients in Group 1 and 2 did not show any fatality and were not included in the survival analysis. In Group 3, the IDH1-WT high-grade glioma group, 37 patients received AcePET-guided surgery and the remaining 43 patients, MRI-guided surgery (Table S1). Multivariate analysis identified AcePET-guided surgery, age, and total resection as independent prognostic factors (Table S2). After propensity score matching (Table S3), we found a significantly prolonged OS in patients with AcePET-guided surgery than with MRI-guided surgery (Figure 7A, Table S4). The AcePET-guided surgery extended the median OS by 5.25 months from 16.75 to 22.00 months and 2-year survival rate by 12.5% from 29.2% to 41.7%. Moreover, the progression-free survival (PFS) was extended by 10.68 months from 8.25 to 18.93 months (Figure 7B, Table S5). These results raise the strong possibility that in IDH1-WT high-grade glioma patients, AcePET-guided surgery provides more accurate tumor boundary information and allows for more complete resection compared to conventional MRI-guided surgery.

**Figure 7.**
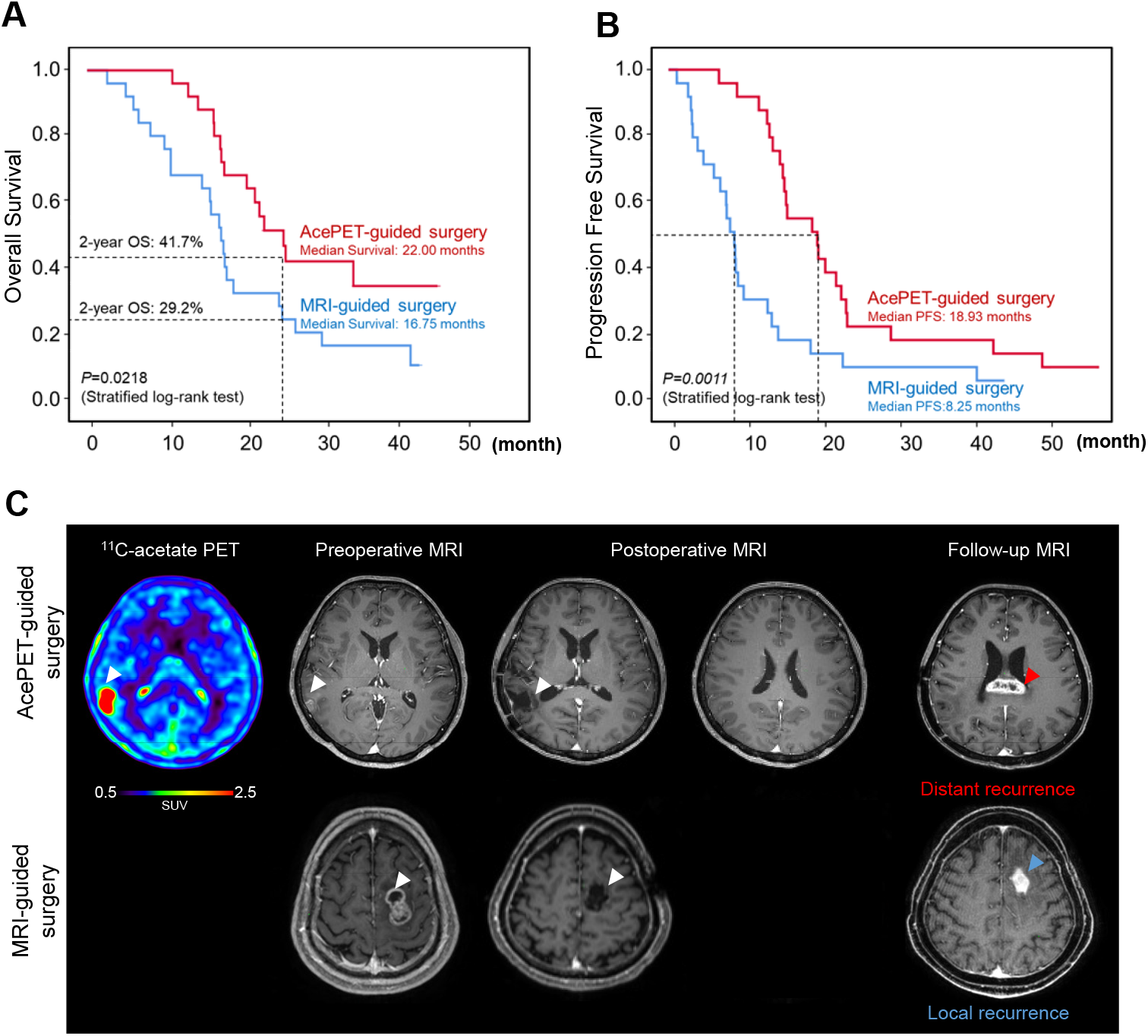
AcePET-guided surgery extends overall survival and reveals reactive astrogliosis at the tumor boundary in high-grade glioma patients. (A) AcePET-guided surgery extended the median OS by 5.25 months from 16.75 to 22.00 months and 2-year survival rate by 12.5% from 29.2% to 41.7%. (B) AcePET-guided surgery extended the median PFS by 10.68 months from 8.25 to 18.93 months. (C) Representative images of ^11^C-acetate PET, MRI in patients with IDH1-WT high-grade gliomas. Upper row: Primary GBM in the right parietal lobe with a total tumor resection on MRI after surgery. Follow-up MRI at 1.5 years postoperatively revealed a recurrence at the corpus callosum. Lower row: Primary GBM in the left superior frontal gyrus with a total tumor resection on MRI after surgery. Follow-up MRI at 9 months postoperatively revealed a recurrence at the resection margin.

To investigate whether AcePET-guided surgery provides less frequent recurrence at the tumor boundary, we further analyzed the 26 patients with total tumor resection on postoperative MRI in the propensity score matching group. Compared to 92% (12 out of 13) of patients with MRI-guided surgery, only 46% (6 out of 13) of patients with AcePET-guided surgery showed tumor recurrence at the resection margin (bottom panel, Figure 7C; P = 0.03). The remaining 56% of patients with AcePET-guided surgery eventually had tumor recurrence at distant regions from the original tumor (top panel, Figure 7C). Taken together, AcePET-guided surgery offers more accurate tumor boundaries, more complete resection, and reduced tumor recurrence at the resection margin, providing a plausible explanation for the extended survival.

## Discussion

One of the most striking findings of our study is that AcePET-guided surgery significantly extended survival in patients with high-grade glioma by 5.25 months of OS and 10.68 months of PFS. This astonishing improvement in patient survival is most likely attributable to more complete removal of cancer stem cells within the regions of reactive astrogliosis beyond the tumor boundary shown on MRI. This improvement was possible due to our newly developed AcePET imaging technology, which is based on acetate uptake by reactive astrocytes. It is the first-of-its-kind to image the TME, whereas other existing imaging techniques, including MRI and 5-ALA fluorescence, are focused on the tumor itself. Cancer stem cells at the leading edge of the tumor play critical roles in rapid expansion and invasion into adjacent normal brain tissue (Croker and Allan, 2008). After surgery, the remaining cancer stem cells are known to be responsible for tumor recurrence (Batlle and Clevers, 2017). In this study, we found less frequent recurrence at the resection margin which clearly provides evidence of better tumor removal by AcePET-guided surgery compared to MRI-guided surgery. Our AcePET-guided surgery provides significant advantages over conventional MRI- or fluorescence-guided surgery, which is constrained by the lack of visualization of reactive astrogliosis and infiltrating cancer stem cells at the leading edges of the tumor (Ji et al., 2015). Despite the prolonged PFS, 54% of patients with AcePET-guided surgery eventually showed tumor recurrence at a site distant from the original tumor. These results are consistent with the recent study reporting the existence of cancer stem cells carrying *p53*, *Pten* and *Egfr* mutations in the subventricular zone (SVZ) (Lee et al., 2018), which could likely be the source of the tumor recurrence in distant brain regions. Therefore, to further improve patient survival in high-grade gliomas, developments in advanced treatment approaches targeting the cancer stem cells in the SVZ are desperately needed.

In this study, we have identified the abnormal multicellular interactions between highgrade glioma cells and reactive astrocytes via excessive acetate, and between reactive astrocytes and neighboring neurons via aberrant GABA and H_2_O_2_ in the TME. The source of such abnormal molecular and cellular communications possibly originates from the monopoly of glucose uptake by high-grade gliomas leading to aggressive tumor proliferation, which is the well-known metabolic hallmark of cancer. Because of the devouring of glucose by highgrade gliomas, the surrounding astrocytes are devoid of available glucose and are forced to use alternative metabolites such as acetate in the TME. While astrocytes under physiological condition utilize glucose to support neuronal metabolism, the increased acetate uptake in reactive astrocytes under GBM synthesize GABA and H_2_O_2_, suppressing neuronal activity in the metabolically challenging TME. The observed metabolic plasticity in reactive astrocytes appears to be at the core of the abnormal multicellular interactions between high-grade glioma cells and neighboring neurons.

The observed metabolic plasticity in reactive astrocytes in GBM is reminiscent of the common metabolic pathway in neurodegenerative diseases such as Alzheimer’s disease, Parkinson’s disease, recovery after stroke&etc. (Heo et al., 2020; Nam, 2021; Nam et al., 2020). The increased acetate in reactive astrocytes synergistically turns on the urea cycle and putrescine metabolism, leading to the aberrant production and release of GABA and H2O2. Consequently, neighboring neurons are strongly suppressed by GABA and H2O2, becoming metabolically poor without glucose. The neuronal hypometabolism and increased inhibitory tone inadvertently cause neuronal dysfunction and cognitive impairment, which are often observed in glioma patients. In our study, we have developed a new way of visualizing reactive astrogliosis and neuronal hypometabolism by using both ^11^C-acetate and ^18^F-FDG PET imaging in GBM patients. The same approach has been successfully applied to patients with Alzheimer’s disease (Nam, 2021). Our approach should be applicable to other neuroinflammatory, infectious, or neurodegenerative diseases, as well as traumatic brain injuries, which all accompany reactive astrogliosis.

We have identified several new molecular targets which can be therapeutically beneficial to anti-cancer treatment, as well as neurodegenerative diseases related to reactive astrogliosis. Firstly, inhibiting acetate release from glycolytic cancer cells may affect tumor growth and stop the cascade of events that lead to reactive astrogliosis and neuronal hypometabolism. Secondly, pharmacological or genetic inhibition of MCT1 should reduce reactive astrogliosis, which may affect tumor growth and invasion. Thirdly, inhibiting MAO-B or ODC1 in reactive astrocytes blocks the MAO-B-dependent putrescine degradation pathway preventing GABA and H2O2 production, which should improve neuronal dysfunction and cognitive impairment. These exciting possibilities await future investigations.

## Acknowledgments

This study was supported by the Institute for Basic Science (IBS-R001-D2) and National Research Foundation (NRF-2018M3C7A1056898, NRF-2020R1A2B5B01098109 and NRF-2021R1C1C2011016) funded by the Korean Ministry of Science and ICT of Republic of South Korea. The schematic figure (Fig. 60) was drawn using Biorender.

## Author Contributions

C.J.L. and M.Y. designed the study. D.K., J.H.C. and M.Y. conceived the idea of AcePET-guided surgery. H.Y.K., D.K., J.C., Y.M.P., Y. P., C.J.L. and M.Y. wrote the manuscript. J.H.C. performed surgeries. D.K managed patient samples and performed statistical analysis in patients. H.Y.K. acquired and analyzed animal images. J.C., H.H.J., S.Y.K., J.K. and M.L. performed and analyzed histological staining. J.C. and H.H.J. performed and analyzed western blot and quantitative RT-PCR. Y.M.P and K.H. performed glutamate and calcium imaging. Y.M.P, K. H. and Y.J performed sniffer patch. Y.M.P and M.H.N. prepared viruses. S.L analyzed PET and MRI images. J.H.C. provided ^11^C-acetate. S.J.P. and K.D.P. provided KDS2010. J.S and S.K. established PDX models. S.H.K. performed pathological studies in patient tissues. All authors contributed to writing and provided feedback.

## STAR methods

### RESOURCE AVAILABILITY

#### Lead contact

Further information and requests for resources and reagents should be directed to and will be fulfilled by the Lead Contact, Mijin Yun.

#### Materials availability

The sequences of the shRNAs used in this study have been provided in the STAR methods. The viruses used in this study were provided by and are available with the Institute for Basic Science Virus Facility (https://www.ibs.re.kr/virusfacility/, Daejeon, South Korea) upon request.

#### Data and code availability

All data reported in this paper will be shared by the lead contact upon reasonable request. No new code has been generated in this study. Any additional information required to reanalyze the data reported in this paper is available from the lead contact upon request.

### EXPERIMENTAL MODEL AND SUBJECT DETAILS

#### Human studies

Between Jan 2016 and October 2017, there were 211 patients with gliomas who had been surgically treated with the aid of image-guided navigation at our institution. Of the 211 patients, 62 were excluded for the following reasons: (a) Follow-up loss (n = 40); (b) Recurrent glioma (n = 17); (c) Patients with double primary cancer (n = 2); (d) Patients who underwent postoperative radiation therapy or concurrent chemoradiation therapy in other hospitals (n = 2); and (e) Pediatric patient (n = 1). There were 149 glioma patients with at least 2 years of follow-up after surgery who received appropriate treatment according to the grade of tumor (Figure S1A). Among them, 95 patients underwent ^11^C-acetate PET/CT as part of a non-randomized prospective clinical trial at Severance hospital. This study was approved by the Institutional Review Board (IRB) of Severance Hospital (4-2011-0697) and all patients provided written informed consent. Also, this study was registered with Clinical Research Information Service of Korea Centers for Disease Control and Prevention (KCT0004468). The study was originally designed to evaluate the value of ^11^C-acetate PET/CT for tumor grading and prediction of patient survival based on the degrees of ^11^C-acetate uptake on PET/CT.

#### Animal

All animal experimental procedures were approved by and performed according to the Institutional Animal Care and Use Committee (IACUC) of Yonsei University College of Medicine. The animals were maintained under maximum barrier specific-pathogen-free conditions with 12 h light-dark cycle at the temperature of 20-22°C and humidity of 35-55%.

##### U87MG or U87mt-orthotopic xenograft

Balb/c nude mice (female; 5-weeks-old; OrientBio) were anesthetized with 2.5% isoflurane and maintained with 2% isoflurane and, before cell implantation, the animal’s head was scrubbed with iodine scrub, and then with alcohol. For implantation, cells were suspended in PBS at a concentration of 1 × 105 cells/μL, and each mouse was stereotactically injected with 2 μL of suspended cells (injection site: AP +0.5; ML −2; DV −3 to bregma). At 2 weeks after postimplantation, the animals were monitored by T2-weighted MRI for the presence of xenographed tumors in the brain. Mice with MRI-identified brain tumors were used for microPET imaging studies.

##### Patient-derived GBM-TS isolation and orthotopic xenograft

Three tumorsphere (TS)-forming GBM cells, TS13-64, TS13-100 and TS19-117 were established from fresh GBM tissue specimens, as approved by the IRB of Severance Hospital (4-2012-0212, 4-2014-0649). TS cells were derived from GBM patients and cultured in TS complete media composed of DMEM/F-12 (Mediatech), 1x B27 (Invitrogen), 20 ng/mL of basic fibroblast growth factor (bFGF, Novoprotein), and 20 ng/mL of epidermal growth factor (EGF, Novoprotein). TS formation followed previous methods (Kong et al., 2013) and the dissociated TS (5 × 10^5^ cells/mL in DMEM/F12) were implanted into the right frontal lobe (injection site: AP +1; ML −2; DV −4.5 to bregma) of Balb/c nude mice (male; 4-8-weeks old; Central Lab. Animal Inc.) using a guide-screw system (Lal et al., 2000). Mice were monitored daily for 120 days after implantation and, if the body weight had decreased by more than 15% compared to their initial weight, were humanely euthanized according to the approved protocol.

#### Cells

##### Glioma cells

U87MG, U373MG, and T98G cancer cells were obtained from Korean Cell Line Bank (KCLB) of Seoul National University. U87-IDH1-mt cells were obtained from American Type Culture Collection (ATCC). U87MG cells were cultured in Dulbecco’s modified Eagle medium (Invitrogen) supplemented with penicillin (100 U/mL; Invitrogen), streptomycin (100 μg/mL; Invitrogen), and fetal bovine serum (FBS) (10% (wt/vol); Sigma-Aldrich) and incubated at 37 °C in a 5% CO2 atmosphere. Cells were used for *in vitro* and *in vivo* experiments when they reached ~80% confluency. All cells were regularly tested for mycoplasma contamination. To prepare the conditioned media (CM), U87MG and U87-IDH1-mt cells were incubated for 1 day (about 80% confluency) and then the culture media were collected and centrifuged to remove cell debris.

##### Primary cells

To isolate primary astrocytes, brain cortices were isolated from ICR mouse (OrientBio) postnatal day 1. The cells from 2 hemispheres were plated on poly-D-lysine (PDL; 100 μg/mL; Sigma)-coated T75 flask in Minimum Essential Media (MEM, Gibco) supplemented with FBS [10% (wt/vol); Hyclone], horse serum [10% (wt/vol); Hyclone], glutamine (2mM; Gibco), penicillin (100 U/mL; Invitrogen), and streptomycin (100 μg/mL; Invitrogen). After 7 days, weakly attached non-astrocytic glial cells were removed by vigorous shaking for 12 h with an orbital shaker (Visionbionex).

Primary neuron-glia co-cultures were obtained from ICR mouse embryos (E14). After brain cortices were isolated and dissociated to single cells through a Pasteur pipette, primary cells were plated at a density of 1 × 10^6^ cells/mL in MEM (Gibco) supplemented with FBS [5% (wt/vol); Hyclone], horse serum [5% (wt/vol); Hyclone], glucose (20 mM), and glutamine (2 mM; Gibco). Cell culture plates were coated with PDL (100 μg/ml; Sigma) and laminin (4 μg/mL; Invitrogen).

Primary neuron cultures were obtained from ICR mouse embryos (E14). After brain cortices were isolated and dissociated to single cells through a Pasteur pipette, primary cells were plated at a density of 1 × 10^6^ cells/mL in Neurobasal A medium (Gibco) supplemented with B27 [2% (wt/vol); Gibco], 1% Glutamax [1% (wt/vol); Thermo], penicillin (100 U/mL; Invitrogen), and streptomycin (100 μg/mL; Invitrogen)Cell culture plates were coated with poly-L-Ornithine hydrobromide (100 μg/ml; Sigma).

## METHOD DETAILS

### PET imaging and image analysis

All patients fasted for at least 6 h before undergoing ^11^C–acetate or ^18^F-FDG injection. For ^11^C–acetate PET/CT, a dose of 740 MBq (20 mCi) of ^11^C–acetate was administered to each patient intravenously and a 20-min emission scan was performed at 10 min after injection. For ^18^F-FDG PET/CT, 4.1 MBq (0.11 mCi) per body weight (kg) of ^18^F-FDG was intravenously administered to each patient and a 15-min emission scan was performed at 60 min after injection. Imaging of the brain was performed on a PET/CT scanner (Discovery 710, General Electric Medical Systems) and PET images were reconstructed using an iterative algorithm (4 iterations, 32 subsets). For attenuation correction, CT scan was obtained with a 0.5 s/rotation, 200 mA, 120 kVp, 3.75-mm section thickness, 10-mm of collimation, and 9.375-mm of table feed per rotation. All PET/CT and MRI images were reviewed and analyzed by one double board-certified radiologist and nuclear medicine physician and one nuclear medicine physician using a fusion module in imaging software (MIM-6.5; MIM Software Inc.) with a decision reached by consensus. The maximum standardized uptake value (SUVmax) of a glioma on ^11^C-acetate PET/CT was measured using a volume of interest (VOI) over the tumor. The contralateral choroid plexus was used as the reference region (Kim et al., 2020). Then, SUV ratio (SUVR) of tumor to choroid plexus was determined as the SUVmax of tumors to the SUVmean of choroid plexus. Tumor volume on ^11^C–acetate PET/CT was calculated using an iso-contour VOI determined using a threshold (SUV_mean_ of the contralateral choroid plexus +2 standard deviation) (Kim et al., 2018).

### MRI imaging and volume measurement

Anatomical datasets were obtained on a 3T whole-body MR scanner (Magnetom Trio, Siemens Medical Solutions) using a contrast-enhanced three-dimensional (3D) T1-weighted magnetization-prepared rapid-gradient-echo (MP-RAGE) sequence. The parameters were as follows: TR=2300 ms; TE=2.98 ms; inversion time=900 ms; flip angle=9°; FOV=256×256 mm; 184×256 matrix with 1×1×1 mm spatial resolution; 224 coronal slices and a slice thickness of 1 mm for 3D T1-weighted MP-RAGE sequences. For tumor volume measurement, tumor margins were manually delineated on the contrast-enhanced axial images over the tumor and all the regions of interest were integrated to measure tumor volume (Roh et al., 2017).

### Image transfer and fusion, and intraoperative image navigation

^11^C–acetate PET and MRI images for navigation surgery were sent from the picture archiving and communication system into the planning station via the local network. A navigation system (StealthViz, Medtronic) was used for the image fusion to guide the surgery. For each patient’s registration, five or more adhesive fiducial skin markers were placed in a scattered pattern on the head before image acquisition, and we registered those markers with a pointer after their position was defined in the 3D dataset. The position of the navigation pointer tip in the surgical field was displayed on the workstation’s monitor with the corresponding location in the image. During resection of the tumor, the neurosurgeon frequently checked anatomical landmarks to ensure accuracy. The tumor margin by AcePET-guided surgery was determined from the fusion images of MRI and ^11^C–acetate PET, and the tumor margin by MRI-guided surgery was based on MRI images in patients without ^11^C–acetate PET. During the surgery, the neurosurgeon performed a targeted biopsy in the regions of high ^11^C–acetate uptake for high-grade gliomas at the tumor boundary and in the peri-tumoral regions for low-grade or IDH1-mt gliomas. Then, the targeted biopsied tissues were transferred to a laboratory for immunohistochemical analysis.

### Tissue preperation for immunostaining of targeted biopsied gloma tissue

For immunostaining of human glioma tissues from the targeted biopsy, tissues were fixed with 4% PFA for 1 day and then made into paraffin blocks and cut into 2-μm sections. After antigen retrieval, the slides were blocked with a blocking solution and incubated with primary antibodies. Primary antibodies were as follows: Rabbit anti-MCT-1 (Alomone, 1:200), mouse anti-MAO-B (Santacruz, 1:200), mouse anti-Ki67 (Dako, 1:500), rabbit anti-GFAP (Dako, 1:500), chicken anti-GFAP (Millipore, 1:500), rabbit anti-CD133 (abcam, 1:200), mouse anti-S100β (Sigma aldrich, 1:400), guinea pig anti-NeuN (Millipore, 1:500), rabbit anti-GLUT3 (Alomone, 1:200), guinea pig anti-GABA (Millipore, 1:200), rabbit anti-GLUT1(abcam, 1:200). After washing in PBS three times, sections were incubated with corresponding fluorescent secondary antibodies (Invitrogen) for 2 hours at room temperature. Secondary antibodies were as follows: Goat anti-chicken-405 (abcam, 1:200), goat anti-chicken-488 (Invitrogen, 1:200), goat anti-chicken-647 (Invitrogen, 1:200), goat anti-guinea pig-488 (Invitrogen, 1:200), goat anti-mouse-488 (Invitrogen, 1:200), goat anti-mouse-594 (Invitrogen, 1:200), goat-anti-mouse-647 (Invitrogen, 1:200), goat anti-rabbit-488 (Invitrogen, 1:200), goat anti-rabbit-594 (Invitrogen, 1:200), goat-anti-rabbit-647 (Invitrogen, 1:200). Following several washes with PBS and counterstaining with Hoechst34580 (Molecular probe, 1:1000), tissue slices were mounted in a fluorescent mounting medium (Dako). A series of fluorescent images were obtained with LSM710 confocal microscope (Carl Zeiss) and 30 μm Z-stack images in 5-μm steps were processed for further analysis using ZEN software (Carl Zeiss).

### Glioma cell line culture

U87MG, U373MG, and T98G cancer cells were obtained from the Korean Cell Line Bank (KCLB) of Seoul National University. U87-IDH1-mt cells were obtained from the American Type Culture Collection (ATCC). U87MG cells were cultured in Dulbecco’s modified Eagle medium (Invitrogen) supplemented with penicillin (100 U/mL; Invitrogen), streptomycin (100 μg/mL; Invitrogen), and fetal bovine serum (FBS) (10% (wt/vol); Sigma-Aldrich) and incubated at 37 °C in a 5% CO2 atmosphere. Cells were used for *in vitro* and *in vivo* experiments when they reached ~80% confluency. All cells were regularly tested for mycoplasma contamination. To prepare the conditioned media (CM), U87MG and U87-IDH1-mt cells were incubated for 1 day (about 80% confluency) and then the culture media were collected and centrifuged to remove cell debris.

### KDS2010 administration

At 1 week after stereotactic injection, tumor formation was confirmed by T2-weighted MRI. Mice were randomly divided into two groups and allowed to drink water (control) or KDS2010 (10 mg/kg daily) containing water at ad libitum. KDS2010, a recently developed selective reversible MAO-B inhibitor, was synthesized as previously described (Park et al., 2019).

### Virus injection

For astrocyte-specific gene-silencing of MCT1, we injected AAV-GFAP-Cre-mCherry (0.5 μL) and AAV-DJ vector containing pSico-MCT1-shRNA-GFP cassette (or pSico-scrambled-shRNA-GFP for control) (0.5 μL) into stereotaxic coordinates of AP +0.5; ML −2; DV −3.2 and −2.8 to bregma using stereotaxic apparatus with a rate of 0.15 mL/min. After injection, the needle was left in place for an additional 7 min and then slowly retracted. The complementary oligomers of mouse MCT1-shRNA were 5’-TGC TCC ACT TAA TCA GGC TTT CTT CAA GAG AGA AAG CCT GAT TAA GTG GAG CTT TTT TC-3’ and 5’-TCG AGA AAA AAG CTC CAC TTA ATC AGG CTT TCT CTC TTG AAG AAA GCC TGA TTA AGT GGA GCA-3’. The viral vectors were packaged by the virus facility at Korea Institute of Science and Technology. The minimum number of viral particles was 1.0 × 10^12^ genome copies (GC)/mL. At 1 week after the virus injection, U87MG cells were injected into AP +0.5; ML −2; DV −3 to bregma for the mouse tumor model.

### Animal microPET and animal MRI

All mice (n=5) were imaged by a microPET scanner (Inveon, Siemens Healthcare). For ^11^C-acetate microPET imaging, mice were intravenously administered with ^11^C-acetate (400 μCi / 0.2 mL Saline), and after 20 min post-injection ^11^C-acetate microPET images were acquired for 40 min. After imaging, animals were kept under a heat lamp for recovery from anesthesia and then placed in an isolation room for 24 h to eliminate radiation hazards. The microPET data were reconstructed with 3-dimensional ordered subset expectation maximization (OSEM) with 2 iterations and 18 subsets. Matrix size was 128 × 128 × 159 and the voxel size was 0.776 × 0.776 × 0.796 mm^3^.

MRI of the implanted brain tumor was performed using a 9.4 T preclinical MR (Bruker BioSpec 94/20 USR, Ettlingen; software: ParaVision 5.0) with water-cooled gradient coils (maximum gradient strength 400 mT/m). Mice were kept warm on a heating pad and anesthetized by 2% isoflurane during the acquisition. MRI was based on respiration triggered and fat suppressed T2-weighted rapid acquisition with relaxation enhancement (RARE) sequence. A 2D set of axial images was acquired (TR, 4900 ms; TEeff, 24 ms; pixel size, 208 μm2; slice thickness, 500 μm).

Image analysis was performed on fused microPET and MRI images using PMOD software (PMOD technologies Ltd.). Two ROIs were manually drawn over PET/MRI fusion images: one ROI on the peri-tumoral and the other on the mirror-image contralateral regions to measure SUVR on microPET.

### Autoradiography

Mouse orthotopic xenograft models (4-5 mice per each group) were intravenously injected with ^14^C-acetate (3 μCi, PerkinElmer) in 200 μL saline and perfused with 4% paraformaldehyde (PFA) at 1 h post-injection. The brains were quickly isolated and frozen in optimal cutting temperature compound (OCT, Leica). Frozen mice brains were sectioned at 20 μm thickness using a cryostat microtome and exposed on an imaging plate (BAS-IP SR 2025, Fuji Film) for 2 weeks. The plates were scanned with phosphor-imager (Typhoon FLA 7000, GE Healthcare).

### Animal tissue immunostaining

Mice were deeply anesthetized with 2%isoflurane and perfused transcardially with 10 mL of heparinized normal saline, followed by ice-cold 4% PFA. Excised brains were post-fixed overnight in 4% PFA and immersed in 30% sucrose for 48 h for cryoprotection. After fixation, brains were mounted into an OCT embedding solution (Leica) and frozen at −20° to −80°C. Tissues were cut in a cryostat by 30-μm coronal sections, then stored in free float storage solution (30% glycerol, 30% ethylene glycol, and 10% 0.2 M phosphate buffer, pH 7.2). Sections were processed with three additional washes in PBS, incubated for 1 h in a blocking solution (0.3% Triton X-100 and 5% normal serum in PBS), and then immunostained with a mixture of primary antibodies in a blocking solution at 4°C on a shaker overnight.

#### In vitro ^14^C-Acetate and ^14^C-Deoxyglucose(DG) uptake assay

In all experiments related to *in vitro* ^14^C-acetate or ^14^C-DG uptake, ^14^C-acetate (PerkinElmer) 0.5 μCi/0.5mL in the external solution (150 mM NaCl, 3 mM KCl, 10 mM HEPES, 22 mM Sucrose, 2 mM MgCl_2_, 2 mM CaCl_2_, pH 7.4) or ^14^C-DG (PerkinElmer) 0.2 μCi/0.2mL in 0.5mM glucose DMEM were added into each well. After 20-min incubation, the external solution was removed and the cells were rinsed twice with ice-cold PBS. Then, the cells were lysed with 200 μL of 0.2N NaOH for 2 h at room temperature. The radioactivity was measured by liquid scintillation counter (Tri-Carb, PerkinElmer) and normalized to protein concentration using a BCA protein assay kit (Thermo Fisher Scientific). Three independent experiments were performed in triplicate.

### DCFDA assay

Intracellular reactive oxygen species (ROS) levels were detected using cell-permeable non-florescent probe 2’, 7’-Dichlorofluorescin diacetate (DCFDA; Invitrogen). DCFDA is de-esterified into its fluorescent form after action of intracellular esterases and oxidation by reactive oxygen species within the cell (Ishii et al., 2017). Primary astrocyte culture was seeded onto 96-well plates (SPL) and treated with U87MG-CM for 2 days. On the 2nd day, cells were washed twice with Hanks Buffered Salt Solution (HBSS; Welgene) and incubated with 30μM DCFDA in HBSS at 37°C for 30 min in the dark. The DCFDA was replaced with HBSS and fluorescence was measured using SpectraMax iD5 Multi-Mode Microplate Reader (Molecular Devices; excitation 485nm emission 538nm).

### siRNA transfection

Primary astrocytes were transfected with siRNA by adding 125 μl of Lipofectamine RNAiMAX (Invitrogen) and 50pmol of siRNA (AccuTarget™ Genome-wide Predesigned siRNA, Bioneer) with OPTIMEM medium (Gibco) following the manufacturer’s instructions. Western blotting and ^14^C-acetate uptake were conducted at 2 days after transfection.

### Western blotting

The cells were subsequently lysed with RIPA lysis buffer (Thermo Fisher Scientific) containing protease inhibitor (Roche), and 10 μg of proteins were separated by electrophoresis in an 10% SDS polyacrylamide gel and transferred onto a polyvinylidene fluoride membrane (PVDF, Millipore). The membranes were blocked with 5% (v/v) skim milk in Tris-buffered saline with 0.1% Tween 20 and incubated with the required antibodies. Primary antibodies were as follows: Rabbit anti-GFAP (Dako, 1:2000), rabbit anti-MCT-1 (Alomone, 1:500), rabbit anti-MAO-B (NOVUS, 1:1000), mouse anti-NeuN (Millipore, 1:1000), rabbit anti-Glut3 (Alomone, 1:1000), mouse anti-β-actin (Invitrogen, 1:5000). Secondary antibodies used were as follows: Goat anti-rabbit-HRP (GeneTex, 1:2000), goat anti-rabbit-HRP (GeneTex, 1:2000). Immunoreactive protein bands were visualized via enhanced chemiluminescence and scanned with a LAS-4000 system (GE Healthcare).

### Quantitative real-time RT-PCR

The RNAs were isolated from mouse primary astrocytes, which were treated with U87MG-CM, U87-IDH1-mt-CM and DMEM (control) for 2 days, using TRIzol reagent (Invitrogen). cDNA was synthetized from 1μg of RNA by iScript^™^ cDNA Synthesis kit (BioRad) and quantitatively amplified using SYBR Green Real-time PCR Master Mix (BioRad). GAPDH was used as an endogenous control to standardize the amount of RNA in each reaction. The sequences of primers were as follows: mouse *Mct1* forward: 5’-GGT CTT GGG CTT GCT TTC AA-3’; *Mct1* reverse: 5’-GAG CCA GGG TAG AGA GGA AC-3’; *Gapdh* forward: 5’-ACC CAG AAG ACT GTG GAT GG-3’, *Gapdh* reverse: 5’-TCA GCT CAG GGA TGA CCT TG-3’, *Cps1* forward: 5’-CTG GCT GGC TAC CAA GAG TC-3’, *Cps1* reverse: 5’-ATT GGG ATC CAC AAA ATC CA-3’*Odc1* forward: 5’-CGT CAC TCC CTT TTA CGC AG-3’, *Odc1* reverse: 5’-AGA TAA CCC TCT CTG CAG GC-3’.

### Acetate assay

For the measurement of acetate amount in U87MG-CM, the colorimetric assay was performed using acetate assay kit (Abcam), according to the manufacturer’s recommendations. DMEM was used as a negative control. The conditioned media (CM) from U87MG were collected and centrifuged for 10 min at 1000 rpm and added into each well of 96 well plate, followed by the reaction mixture addition. The plate was incubated at room temperature for 40 min while protected from light. To analyze results, absorbance at 450 nm was measured using a micro-plate reader (Molecular Devices).

### Sniffer patch

The sensor cells were originated from HEK293T cell line with stable expression of GABAc receptor (GABAc). The construction of the stable cell line is described as follows; The full-length of GABAc CDS was cloned into pHR-CMV-IRES-EmGFP lentiviral expression vector (a gift from A. Radu Aricescu, Addgene plasmid # 113888) via AgeI and EcoRI restriction sites. For lentiviral particle production, the cloned plasmids were transiently transfected into HEK293T cells with packaging plasmid psPAX2 and envelope plasmid pMD2.G using Lipofectamine 3000 transfection reagent (Invitrogen) based on the manufacturer’s instructions. At 24 h and 48 h post-transfection, the cell culture media containing lentivirus particles were harvested, filtered through 0.45 μm-pore-size filters, and stored at −80°C until needed. For the generation of stable GABAc-expressing sensor cell line, lentiviral particle-containing media were directly overlaid on HEK293T cells in the presence of polybrene (Merck) at a final concentration of 4 μg/mL. After 24 h, the supernatant was changed with fresh medium and incubated for 2 days. GFP-positive cells were single-cell sorted into 96-well plates using a MoFlo Astrios Cell Sortor (Beckman Coulter) and cultured for another 2~3 weeks to allow for clonal expansion. HEK293T cells were regularly tested for mycoplasma contamination.

The day before sniffer patch, cortical astrocytes were seeded from culture dishes onto 12-mm glass coverslips coated with PDL in 24-well plates. On the day of the sniffer patch, sensor cells were seeded from culture dishes onto astrocyte-placed coverslips. Astrocytes, which were co-cultured with GABAc sensor cells, were incubated with 5 μM Fura-2AM for 40 min and washed at room temperature and subsequently transferred to a microscope stage with external solution. For Ca^2+^ imaging, the intensity of 510 nm wavelength was imaged at 340 nm and 380 nm excitation wavelengths using CoolLED (pE-340fura). Astrocytic Ca^2+^ responses were induced by poking as previously described (Oh et al., 2019). GABAc-mediated currents from sensor cell were simultaneously recorded under voltage clamp (Vh = - 50 mV) using Multiclamp 700B amplifier (Molecular Devices), acquired with pClamp 11.0.3 Recording electrodes (4-7 MΩ) were filled with: 140 mM CsCl, 0.5 mM CaCl_2_, 10 mM HEPES, and 10 mM EGTA (pH adjusted to 7.3 with CsOH and Osmolality with 285–295 mOsm/kg). To normalize the different expression of GABAc in sensor cells, 100 μM of GABA was bath-applied to obtain the maximal GABAc current from each sensor cell. The sniffer-current, mediated by the released GABA from astrocytes, was normalized by the maximal GABAc current.

## QUANTIFICATION AND STATISTICAL ANALYSIS

### Statistical analysis for clinical data

In this study, we retrospectively analyzed patient survival between AcePET-guided surgery and conventional MRI-guided surgery in IDH1-WT high-grade gliomas. We recruited patients who underwent ^11^C–acetate PET/CT from the previous prospective clinical trial (Table S1). Univariate and multivariate Cox proportional hazards regression analyses were used to evaluate the efficacy of prognostic factors including age, sex, Karnofsky Performance Status Scale (KPS), Ki-67, Surgery (Total resection versus Subtotal resection), O-6-methylguanine DNA methyltransferase (MGMT) status, and performance of ^11^C–acetate PET/CT in IDH-WT high-grade glioma patients (Table S2). The relationship between the histological WHO grades (Grade 2 Versus Grade 3, 4) and SUVR on ^11^C–acetate PET was assessed using the Mann-Whitney U-test. The statistical differences in tumor volumes between MRI and PET were also assessed using the Mann-Whitney U-test. All tests were two-sided.

As a retrospective study, we conducted propensity score matching (PSM) to reduce inevitable selection bias due to unbalanced baseline patient characteristics. Briefly, the propensity scores of all IDH1-WT high-grade glioma patients were calculated by a logistic regression model, in which AcePET-guided surgery was considered the dependent variable with regard to all clinicopathological covariates presented in Table S1. Patients who received AcePET-guided surgery were matched to those received MRI-guided surgery according to the propensity scores by the nearest neighbour matching without replacement at 1:1 fixed ratio. After matching, the standardized mean differences in each of the covariates were applied to compare the balance of the matched cohort (Table S3). Univariable stratified cox regression analysis was used to evaluate the efficacy of AcePET-guided surgery for OS and PFS after PSM (Table S4 and S5). Finally, OS and PFS were analyzed using Kaplan-Meier survival curves and stratified log-rank test in IDH1-WT high-glioma patients after PSM. All statistical analyses were performed using IBM SPSS Statistics for Windows, version 25.0 (IBM Corp), and SAS (version 9.4, SAS Inc.). A P-value less than 0.05 was considered statistically significant.

### Image analysis of immunostaining

Immunostained sections were imaged with Zeiss LSM710 confocal microscope. Confocal images were analyzed using the ImageJ program (NIH). For the measurement of the immunoreactivity of MCT1 or MAOB in astrocytes, we first generated binary images after setting the threshold of the GFAP-positive pixels (GFAP+) or S100β+ to define single astrocytes as regions-of-interest (ROIs). Then the intensity of MCT1 or MAOB in each GFAP+ or S100β+ ROI was measured. For the measurement of GLUT3 levels in neurons, we generated the binary NeuN+ images to define single neurons as ROIs. Then we measured the intensity of GLUT3 in each NeuN+ ROI. For the measurement of GFAP+ or S100β+ area, we analyzed the area of each GFAP+ or S100β+ ROI. For morphological analysis of astrocytes, we utilized the Surface tool of Imaris 9 (Bitplane) for 3-D rendering GFAP+ cells, and then conducted the Sholl analysis in ImageJ.

### Statistial Analysis for in vitro and in vivo animal studies

All data were presented as mean ± SEM. Statistical analyses were performed using Prism 8 (GraphPad Software, Inc.). The normality test was conducted on each dataset. If the data were not distributed normally, non-parametric tests were adopted. To determine the statistical significance, P values were calculated using two-tailed Student’s unpaired t-test, one-way analysis of variance (ANOVA) with Tukey’s multiple comparison test, Mann-Whitney U tests, or Kruskal-Wallis test. The significance level is represented as asterisks (*P < 0.05, **P < 0.01, ***P < 0.001; ns, not significant). The number of animals used is described in the corresponding figure legends or on each graph. All experiments were done with at least three biological replicates. Experimental groups were balanced in terms of animal age, sex and weight. Prior to virus injection or drug administration, animals were randomly and evenly allocated to each experimental group. However, investigators were not blinded to group allocation during experiments or to outcome assessments.

## Supplemental Information

**Figure S1**. Sholl analysis and orientation plot of reactive astrocytes. (A) Representative image for Sholl analysis of an astrocyte according to S, L or T region in the human glioma tissues. (B) Orientation plot of S and L astrocytes in Figure 2c (related to Figure 2).

**Figure S2**. Reactive astrocytes showed high GFAP, MCT1 and MAOB expression. (A) Immunofluorescence images with GFAP. (B) Immunofluorescence images with of S100β and MCT1 (left) or GFAP and MAO-B (right). (C) Immunofluorescence images with GFAP. (D) Immunofluorescence images of S100β and MCT1 (left), GFAP and MAO-B (right). (related to Figure 3)

**Figure S3.** (A) Procedure of AcePET-guided surgery. (B) Flow chart of the study population. (related to Figure 7)

**Table S1.**Patient demographics and histopathological characteristics. (related to Figure 7 and STAR methods)

**Table S2**. Univariate and multivariate Cox proportional hazards analyses for overall survival in IDH1-wildtype high-grade gliomas. (related to Figure 7 and STAR methods)

**Table S3**. Patient characteristics before and after propensity score matching. (related to Figure 7 and STAR methods)

**Table S4**. Univariable stratified cox regression results in overall survival. (related to Figure 7 and STAR methods)

**Table S5**. Univariable stratified cox regression results in progression free survival. (related to Figure 7 and STAR methods)

