## Supplementary information for "Visualizing cancer-originated acetate uptake through MCT1 in reactive astrocytes demarcates tumor border and extends survival in glioblastoma patients"

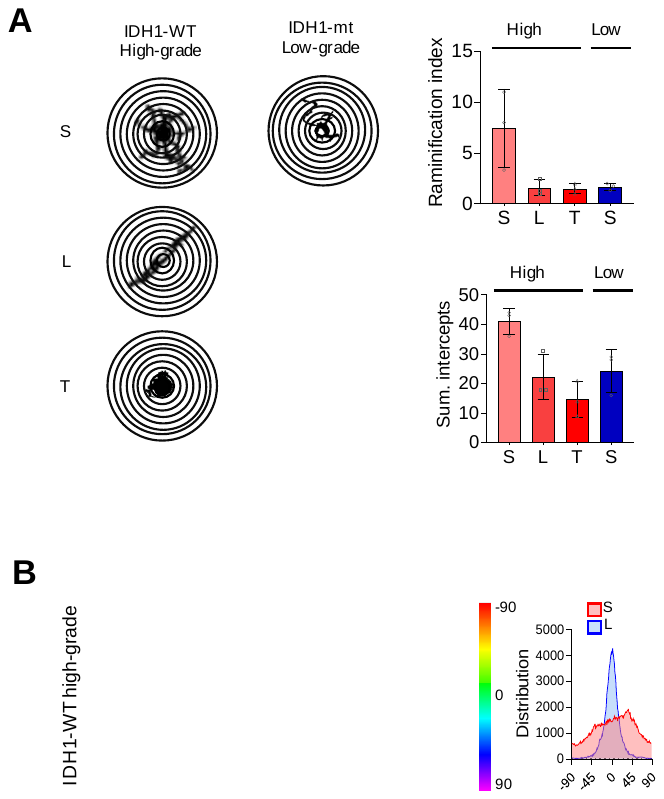

**Figure S1. Sholl analysis and orientation plot of reactive astrocytes, related to Figure 2**. (A) Representative image for Sholl analysis of an astrocyte according to S, L or T region in the human glioma tissues. (B) Orientation plot of S and L astrocytes in Figure 2c,

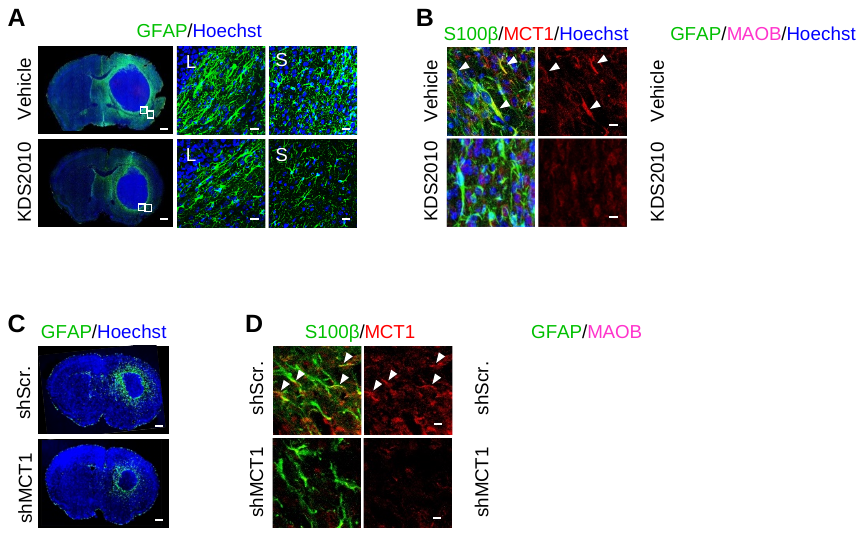

**Figure S2. Reactive astrocytes showed high GFAP, MCT1 and MAOB expression**, **related to Figure 3. (A)** Immunofluorescence images with GFAP. (**B**) Immunofluorescence images with of S100β and MCT1 (left) or GFAP and MAO-B (right). (**C**) Immunofluorescence images with GFAP. **D**, Immunofluorescence images with of S100β and MCT1 (left), GFAP and MAO-B (right).

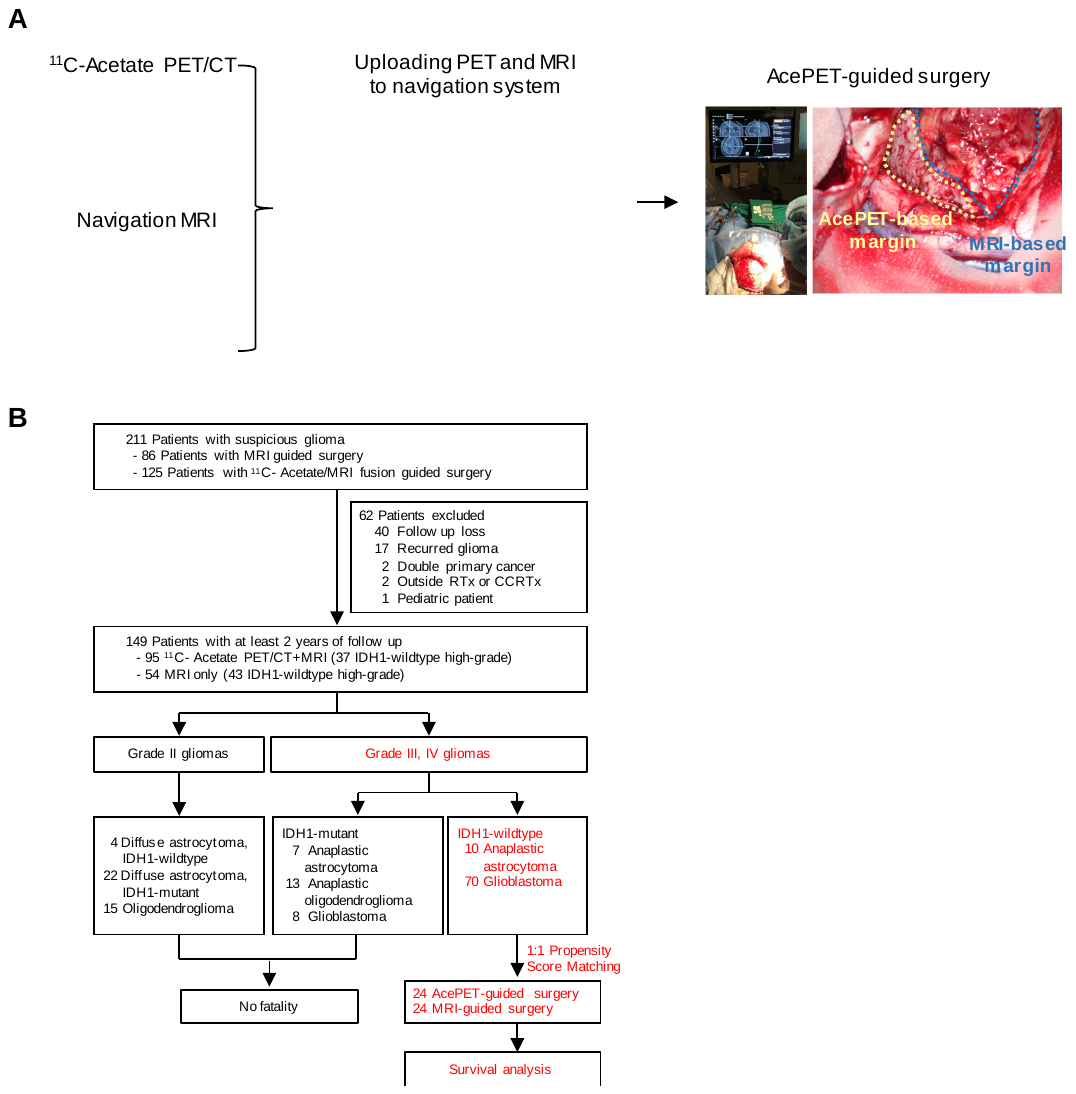

**Figure S3.** (A) Procedure of AcePET-guided surgery (B) Flow chart of the study population (related to Figure 7)

**Table S1. Patient demographics and histopathological characteristics, related to Figure 7 and STAR methods**

| **Characteristic** | | **Values** |
| --- | --- | --- |
| Age (years) | | Median: 59.0 (range, 25-88) |
| Sex, n (%) | Male | 54 (67.5%) |
|  | Female | 26 (32.5%) |
| WHO 2016 grade | Grade Ⅲ | 10 (12.5%) |
|  | Grade Ⅳ | 70 (87.5%) |
| Surgery | Total | 49 (61.3%) |
|  | Subtotal or Partial | 31 (38.7%) |
| Navigation surgery | Acetate/PET-guided | 37 (47.4%) |
|  |  | Grade Ⅲ: 7, Grade Ⅳ: 30 |
|  | MRI-guided | 43 (52.6%) |
|  |  | Grade Ⅲ: 3, Grade Ⅳ: 40 |
| MGMT | Methylated | 33 (41.3%) |
|  | Unmethylated | 47 (58.7%) |
| Ki-67 | | Median: 25.0 |
|  | | (IQR 10.0–40.0) |

**Table S2**. **Univariate and multivariate Cox proportional hazards analyses for overall survival in IDH1-wildtype high-grade gliomas, related to Figure 7 and STAR methods**.

| **Variables** | **Univariate** | |  | | **Multivariate** | |
| --- | --- | --- | --- | --- | --- | --- |
|  | **Hazard ratio (95% CI)** | **P value** | |  | **Hazard ratio (95% CI)** | **P value** |
| Age | 1.026 (1.005-1.049) | 0.017 | |  | 1.025 (1.002-1.049) | 0.032 |
| KPS | 0.982  (0.962-1.003) | 0.087 | |  |  |  |
| Ki-67 | 1.012  (1.000-1.024) | 0.053 | |  |  |  |
| Sex | 0.589  (0.330-1.050) | 0.073 | |  |  |  |
| Surgery | 2.017  (1.188-3.425) | 0.009 | |  | 2.026 (1.173-3.502) | 0.011 |
| MGMT | 1.378  (0.808-2.347) | 0.239 | |  |  |  |
| Acetate | 2.322  (1.351-3.990) | 0.002 | |  | 1.829 (1.037-3.227) | 0.037 |

**Table S3. Patient characteristics before and after propensity score matching, related to Figure 7 and STAR methods**. After PSM was performed using age, sex, surgery, KPS, MGMT, and Ki67, the significance of the variables between the two groups was examined. All variables used in PSM (age, sex, surgery, KPS, MGMT, Ki67) can be considered to be well matched only when there is no significant difference between the two groups. There is no difference, so it can be said that the match was successful.

| **Variables** | **Label** | **Before matching** | | |  |  | **1:1 PSM** | |  |
| --- | --- | --- | --- | --- | --- | --- | --- | --- | --- |
|  |  | **0 (No acetate)**  **(N=43)** | **1 (Acetate)**  **(N=37)** | **p-value** |  | **0 (No acetate) (N=24)** | | **1 (Acetate)**  **(N=24)** | **p-value** |
| Age |  | 62.12±13.18 | 55.38±13.25 | 0.0070 |  | 58.75±13.50 | | 58.79±10.40 | 0.9904 |
| KPS |  | 80 (70 , 90) | 80 (80 , 100) | 0.0288 |  | 90 (80 , 90) | | 80 (80 , 90) | 0.6547 |
| Ki-67 |  | 30 (10 , 50) | 20 (8 , 30) | 0.0164 |  | 20 (10 , 35) | | 20 (10 , 30) | 0.9093 |
| Sex | 0: Male | 30 (69.77) | 25 (67.57) | 0.8324 |  | 15 (62.50) | | 16 (66.67) | 0.7815 |
|  | 1: Female | 13 (30.23) | 12 (32.43) |  |  | 9 (37.50) | | 8 (33.33) |  |
| Surgery | 0: Total | 24 (55.81) | 25 (67.57) | 0.2820 |  | 14 (58.33) | | 15 (62.50) | 0.7389 |
|  | 1: Sub | 19 (44.19) | 12 (32.43) |  |  | 10 (41.67) | | 9 (37.50) |  |
| MGMT |  | 20 (46.51) | 13 (35.14) | 0.3027 |  | 9 (37.50) | | 8 (33.33) | 0.6547 |
|  |  | 23 (53.49) | 24 (64.86) |  |  | 15 (62.50) | | 16 (66.67) |  |
| Event | 0: Survival | 6 (13.95) | 16 (43.24) | 0.0034 |  | 4 (16.67) | | 9 (37.50) | 0.1317 |
|  | 1: Death | 37 (86.05) | 21 (56.76) |  |  | 20 (83.33) | | 15 (62.50) |  |

**Table S4. Univariable stratified cox regression results in overall survival,** **related to Figure 7 and STAR methods.** In the 1:1 PSM results, the mortality rate was significantly lower in the group receiving AcePET-guided surgery compared to the other group when using PSM. It can be said that the risk of death in the group receiving AcePET-guided surgery is significantly lower by 0.524 times compared to the other group

| **Variables** | **Label** | **1:1 PSM** | | | |
| --- | --- | --- | --- | --- | --- |
|  |  | HR | Lower | Upper | p-value |
| Acetate | 0: No acetate | 1 (ref) |  |  |  |
|  | 1: Acetate | 0.457 | 0.255 | 0.820 | 0.0086 |

**Table S5. Univariable stratified cox regression results in progression free survival, related to Figure 1 and STAR methods.** In the 1:1 PSM results, the risk of progression in the group receiving AcePET-guided surgery is significantly lower by 0.377 times compared to the other group.

|  |  | **HR (95% CI)** | **p-value** |
| --- | --- | --- | --- |
| Acetate | 0: No acetate | 1 (ref) |  |
|  | 1: Acetate | 0.377 (0.217 - 0.654) | 0.0005 |
